# High-throughput glycan array screening reveals rhamnogalacturonan-I as a ligand for Arabidopsis leucine-rich repeat receptor kinases

**DOI:** 10.1101/2025.01.29.635407

**Authors:** Du-Hwa Lee, Colin Ruprecht, Jung-Min Lee, Min-Soo Choi, Tobias Hrovat, Hyeonmin Ryu, Sejin Choi, Geon Heo, Natalie Edelbacher, Balaji Enugutti, Markus Blaukopf, Chung Hyun Cho, Ho-Seok Lee, Youssef Belkhadir, Elwira Smakowska-Luzan, Fabian Pfrengle

**Author notes:** These authors contributed equally (Du-Hwa Lee, Colin Ruprecht).

## Abstract

The plant cell wall not only serves as a physical barrier against pathogens but, when damaged, also functions as a source of cell wall-derived molecules that play crucial roles in plant immunity as damage-associated molecular patterns (DAMPs). While oligogalacturonides from homogalacturonan are well-studied DAMPs, the immune-signaling potential of other cell wall components remains largely unexplored. Conventional genetic and biochemical approaches aimed at identifying ligand-receptor pairs in plant immunity have been limited by the vast diversity of potential ligand molecules and functional redundancy of putative receptors. Here, we developed a high-throughput screening pipeline that simultaneously examines multiple interactions between plant cell wall-derived glycans and >350 extracellular domains (ECDs) of receptor kinases and receptor like proteins in Arabidopsis, resulting in the screening of >40,000 interactions. We discovered a group of leucine-rich repeat receptor kinases named ARMs (AWARENESS of RG-I MAINTENANCES) that interact with rhamnogalacturonan-I (RG-I), a major component of pectin. RG-I treatment induced pattern-triggered immunity responses, with distinct kinetics compared to oligogalacturonide responses. We identified RG-I oligosaccharide structures required for interaction with ARM receptors and immune activation, and found that ARM receptors are redundantly involved in plant immunity. Collectively, our approach provides a powerful platform for discovering glycan-receptor pairs in plants, facilitating a more comprehensive understanding of cell wall surveillance mechanisms in plant immunity.

## Introduction

The cell wall, as the outermost structure, serves as a primary barrier for plant cells against interacting organisms (Wolf, 2022). The plant cell wall contains large amounts of the polysaccharides, including cellulose, hemicellulose, and pectin, which provide structural integrity and support (Delmer et al., 2024). Due to its outermost nature, the cell wall is the first target to break down during pathogen or pest attacks (Malinovsky et al., 2014). In the context of innate immunity, products of cell wall breakdown can be recognized as typical extracellular warning signals (Molina et al., 2024). Cell wall components of attacking organisms are categorized as microbe-associated molecular patterns (MAMPs), whereas host cell wall components produced by mechanical forces or digestion of enzymes by attacking organisms are categorized as damage-associated molecular patterns (DAMPs) (Bacete et al., 2018). Plants have evolved an expanded repertoire of cell surface receptors to respond to such MAMPs/DAMPs (Lee et al., 2021). This subset of cell surface receptors is called pattern recognition receptors (PRRs), triggering pattern-triggered immunity (PTI) upon binding these molecular patterns (DeFalco and Zipfel, 2021). For example, PRRs of the lysin motif (LysM) family, CHITIN ELICITOR RECEPTOR KINASE 1 (CERK1) and LysM motif receptor kinase 4 and 5 (LYK4 and LYK5) can recognize the glycan-type MAMP chitohexaose (β-1,4-D-N-acetylglucosamine)_6_ from fungal pathogens (Cao et al., 2014; Kaku et al., 2006; Miya et al., 2007). To date, some glycan-type DAMP ligand-receptor pairs have been extensively studied: firstly, Arabidopsis wall-associated kinases 1 and 2 (WAK1 and WAK2) have been shown to interact with the glycan-type DAMP oligogalacturonic acid (⍺-1,4-oligogalacturonan; OGs) derived from plant pectin, and have been implicated in OGs perception (Brutus et al., 2010; Hématy et al., 2009). Among OG fragments, those with 10-15 degrees of polymerization (OG^DP10-15^) have been shown to induce the strongest immune responses (Côté and Hahn, 1994; Denoux et al., 2008; Xiao et al., 2024). However, two independent recent studies using a quintuple deletion mutant of WAK1 and its homologs WAK2-5, generated by the clustered regularly interspaced short palindromic repeats (CRISPR)/CRISPR-associated 9 (Cas9) technique, showed no or only minor changes in PTI response to OGs (Herold et al., 2024; Kohorn et al., 2021), suggesting functional redundancy between WAK receptors with WAK-like (WAKL)-family receptors. More recently, PRRs containing leucine-rich-repeat (LRR) and malectin (MAL) domains, namely CELLO-OLIGOMER RECEPTOR KINASE1/IMPAIRED IN GLYCAN PERCEPTION 1, 2, and 4 (CORK1/IGP1-4), have been identified as receptors for cello-oligomers (β-1,4-oligoglucans), such as cellotriose (Martín-Dacal et al., 2022; Tseng et al., 2022). These studies revealed direct binding between CORK1/IGP receptors and cello-oligosaccharides resulting in impaired immune responses in receptor mutants. Given the diversity of glycan compositions in the plant cell wall and the wide range of cell wall-degrading enzymes (CWDEs) secreted by pathogens, an array of cell wall derivatives can be produced as MAMPs/DAMPs during pathogen attack (Delmer *et al*., 2024; Kubicek et al., 2014). Indeed, a significant number of glycans have been identified to trigger PTI responses, including (arabino-)xylan, mixed linkage glucan (MLG), xyloglucan and mannan oligosaccharides (Aziz et al., 2007; Bacete et al., 2017; Claverie et al., 2018; Klarzynski et al., 2000; Molina *et al*., 2024; Mélida et al., 2020; Mélida et al., 2018; Wanke et al., 2020). While putative receptors for some of these glycan ligands have been reported, substantial opportunity remains for discovering novel glycan ligands that trigger PTI responses as well as investigating their ligand specificities and the molecular mechanisms resulting upon ligand perception.

The general responses of PTIs, even when activated by different ligands, share key conceptual similarities (DeFalco and Zipfel, 2021). Upon detecting MAMPs/DAMPs, the corresponding receptor kinases (RKs) or associated RKs are activated and initiate downstream signal transduction pathways. Key features of PTI include the activation of ion channels, production of reactive oxygen intermediates (ROIs), activation of defense-related mitogen-activated protein kinases (MAPKs) cascades, and extensive transcriptomic changes (Bigeard et al., 2015; Bjornson et al., 2021). Prolonged MAMPs/DAMPs exposure leads to long-term trade-offs between resource allocation for growth and immune responses, ultimately leading to growth inhibition (Gómez-Gómez et al., 1999; He et al., 2022; Jing et al., 2019). To date, various approaches have been employed to identify receptor-ligand relationships. Forward genetic screens using chemical mutagens such as ethyl methanesulfonate (EMS) have been one of the most successful strategies, relying on monitoring PTI response readouts after known MAMP/DAMP molecules treatment (Choi et al., 2014; Martín-Dacal *et al*., 2022; Ranf et al., 2015; Tseng *et al*., 2022). However, many PRRs have been reported to have redundant functions (Liu et al., 2023; Wang et al., 2018; Yamaguchi et al., 2010), demonstrating the limitations of these forward genetic approaches in determining multiple receptor-ligand relationships.

In this study, we developed a high-throughput screening pipeline to examine the multiple interactions between cell wall-derived glycans and a protein expression library of extracellular domains (ECDs) from RKs and receptor-like proteins (RLPs) in Arabidopsis (Fig. 1A). Using this pipeline, we identified five ECDs of RKs that interact with rhamnogalacturonan-I (RG-I), a component of pectin. These receptors, named AWARENESS of RG-I MAINTENANCES (ARMs), bind RG-I polysaccharides as well as defined RG-I oligosaccharides either purified from natural sources or synthesized and are required for resistance to pathogenic bacterial infection.

**Figure 1.**
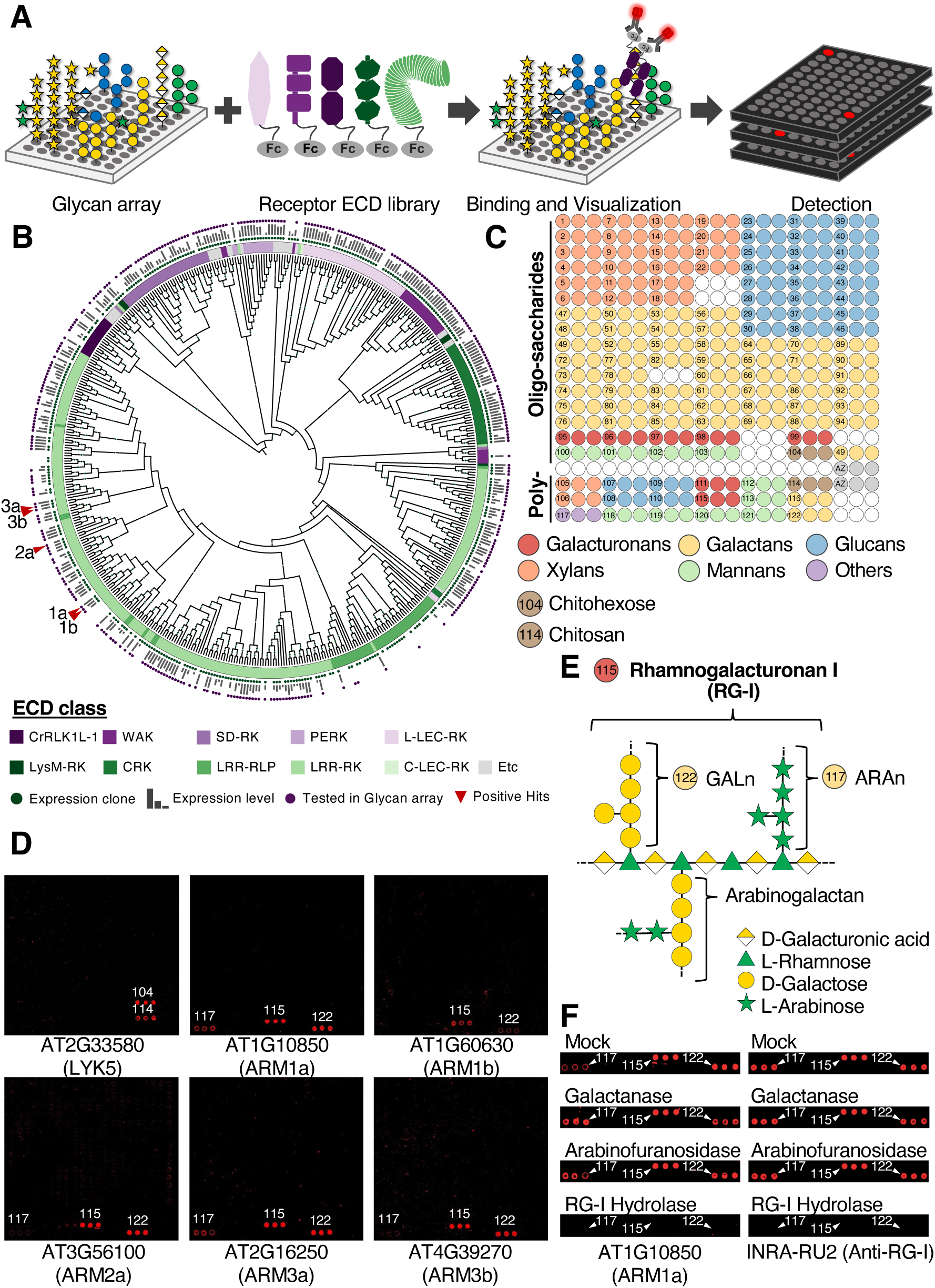
High-throughput plant glycan array screening revealed that five ECDs of LRR-RKs interact with polysaccharides related to Rhamnogalacturonan-I (RG-I). A. A schematic representation of the screening pipeline using a recombinant ECD library of Arabidopsis receptors and a plant glycan array. B. A receptor tree based on ECD sequence similarity of all Arabidopsis plasma membrane-located receptor proteins. ECD classes were distinguished by different color codes. Bootstrap values above 50 are represented with different dot sizes on each node. Detailed information is available in Supplemental Fig. 1. C. Printing pattern for the plant glycan array. Detailed information is available in Supplemental Fig. 3. D. Positive hits from screening results. The chitin receptor LYK5 (AT2G33580) was used as a positive control. Numbers indicate the printed regions that described in (C). E. Proposed structure of RG-I and related polysaccharides that bind to ARM receptors. Numbers in circles indicate glycan array positions: #115 Rhamnogalacturonan I (RG-I; Megazyme P-RHAM1, [4)-α-D-GalA-(1,2)-α-L-Rha-(1]_n_ backbone with β-1,4-galactan and α-1,5-arabinan side chains), #117 Arabinan (ARAn; Megazyme P-ARAB, α-1,5-linked L-arabinofuranose), and #122 Galactan (GALn; Megazyme P-GALPOT, β-1,4-linked D-galactopyranose), corresponding to positions shown in Supplemental Fig. 3. F. On-array digestion was performed with galactanase, arabinofuranosidase, or RG-I hydrolase. ARM1a^ECD^ was used to determine binding on the glycan array after digestion. The RG-I-specific antibody INRA-RU2 (anti-RG-I backbone) was used to confirm the clear digestion. Numbers indicate the printed regions that described in (C).

## Results

We previously established an Arabidopsis LRR-RK network using about 200 ECDs in a recombinant protein expression library for *Drosophila* Schneider S2 cells (Smakowska-Luzan et al., 2018). We have since expanded this library to encompass 422 ECDs from RKs and RLPs in the Arabidopsis genome, covering a broader range of receptor classes (Fig. 1B and Supplemental Fig. 1). 357 of these proteins were subsequently used in binding assays with a glycan array populated with 104 synthetic plant oligosaccharides and 18 commercially available plant polysaccharides (122 total), resulting in the screening of 43,554 protein-glycan combinations (Fig. 1C). Expression levels and estimated sizes for these proteins were evaluated by immunoblotting assays (Supplemental Fig. 2 and Supplemental Table 1). The chemically synthesized oligosaccharides include arabinoxylan, type I and type II arabinogalactans, xyloglucan, rhamnogalacturonan-I, and mixed-linkage glucan oligosaccharides, representing fragments of natural hemicelluloses and pectins, as well as hydroxyproline-rich glycoproteins (Supplemental Fig. 3) (Ruprecht et al., 2017; Ruprecht et al., 2020). To ensure that receptor-glycan interactions can be detected using the glycan array assay, we performed proof-of-concept binding studies using LYK5^ECD^ and CERK1^ECD^. Both proteins bound to chitohexaose and chitosan polysaccharides on the array, with LYK5^ECD^ interacting stronger, particularly to chitosan compared to CERK1^ECD^ (Fig. 1D, Supplemental Fig. 4A, and Supplemental Table 2). Besides LYK5^ECD^ and CERK1^ECD^, which served as stringent and relaxed thresholds respectively, 25 additional hits showed fluorescence signals above the CERK1^ECD^ threshold. Among these hits, 12 hits from four ECDs were determined to be false positive signals because these signals resulted from technical artifacts (Supplemental Fig. 4B and Supplemental Table 2). Thus, the remaining 13 hits from five ECDs from a subfamily of LRR-RKs represent genuine binding results with glycans on the array. All five ECDs bound to the polysaccharides rhamnogalacturonan-I (RG-I, #115), with some also binding to arabinan (ARAn, #117) and galactan (GALn, #122) (Fig. 1D and Supplemental Table 2). The backbone of RG-I consists of alternating units of D-galacturonic acid (GalA) and L-rhamnose (Rha) [4)-α-D-GalA-(1,2)-α-L-Rha-(1]_n_, with the Rha residues partially being substituted by side chains such as galactan (GALn; β-1,4-galactan), arabinan (ARAn; α-1,5-arabinan), and arabinogalactan (β-1,4-galactan with α-1,5-arabinan branches) (Fig. 1E). While ARAn and GALn can exist as independent polysaccharides, the commercially available ARAn and GALn polysaccharides used in our array were originally purified from plant pectin and might thus contain RG-I contamination. To confirm the presence of RG-I in the GALn and ARAn samples, we used the monoclonal antibody INRA-RU2, which specifically binds to the RG-I backbone (Ralet et al., 2010; Ruprecht *et al*., 2017). INRA-RU2 bound in addition to RG-I also to the GALn and ARAn samples on the array, indicating the presence of RG-I in these samples (Fig. 1F). To determine if these ECDs bind to the RG-I backbone, we selected one positive hit (AT1G10850) and performed on-array digestion using an endo-RG-I hydrolase. We found that the binding of this ECD was abolished when RG-I was digested, indicating that the RG-I backbone structure is essential for binding. We therefore named these five receptors AWARENESS of RG-I MAINTENANCES (ARMs).

Based on ECD protein sequence similarity analysis, the five ARM receptors were distributed across 3 different clades, which we named ARM1, ARM2, and ARM3 with increasing numbers of LRRs and ECD sizes, respectively (Fig. 2A and Supplemental Fig. 5A and B). Within these clades, we found pairs of positive hits that were closest homologs to each other in the ARM1 and ARM3 clades, which we designated as ARM1a (AT1G10850)/ARM1b (AT1G60630) and ARM3a (AT2G16250)/ARM3b (AT4G39270), respectively. The ARM2 clade contained only one positive hit, AT3G56100, which we designated as ARM2a. Although its closest homolog AT3G51740 did not show positive interactions in our screening (Supplemental Fig. 5C), we designated it as ARM2b and included it in further detailed interaction assays. To quantify the interaction between ARM^ECD^s and RG-I polysaccharides, we purified the ARM^ECD^s and LYK5^ECD^ (Supplemental Fig. 6A) and performed microscale thermophoresis (MST) assays. Due to the heterogeneous nature of polysaccharides, we determined the half-maximal effective concentration (EC50) values rather than dissociation constants (Kd). The five ARM^ECD^s that showed positive interactions with RG-I polysaccharides in the screening exhibited EC50 values ranging from 31.85±6.23 µg/mL (ARM2a^ECD^) to 70.27±14.09 µg/mL (ARM1b^ECD^) (Fig. 2B). Interestingly, while ARM2b^ECD^ did not bind to RG-I on the array, it showed interaction with RG-I in MST, albeit with a higher EC50 value of 119.2±22.84 µg/mL (Fig. 2B), indicating a weaker interaction compared to the other ARM^ECD^s. This result suggests that the sensitivity limit in the glycan array within RG-I interactions corresponds to EC50 values between ARM2b^ECD^ and ARM1b^ECD^, i. e. 70-120 µg/mL. As a positive control, binding between LYK5^ECD^ and chitopentaose (GlcNAc)_5_ was confirmed with a Kd of 9.01 ± 4.41 µM (Fig. 2C). Additionally, we were unable to detect interactions between ARM1a^ECD^ and OG^DP10-15^ or between a randomly selected large LRR^ECD^ (AT5G14210) and RG-I in the MST assay (Fig. 2C and D), indicating the specificity of the interactions between ARM^ECD^s and RG-I polysaccharides.

**Figure 2.**
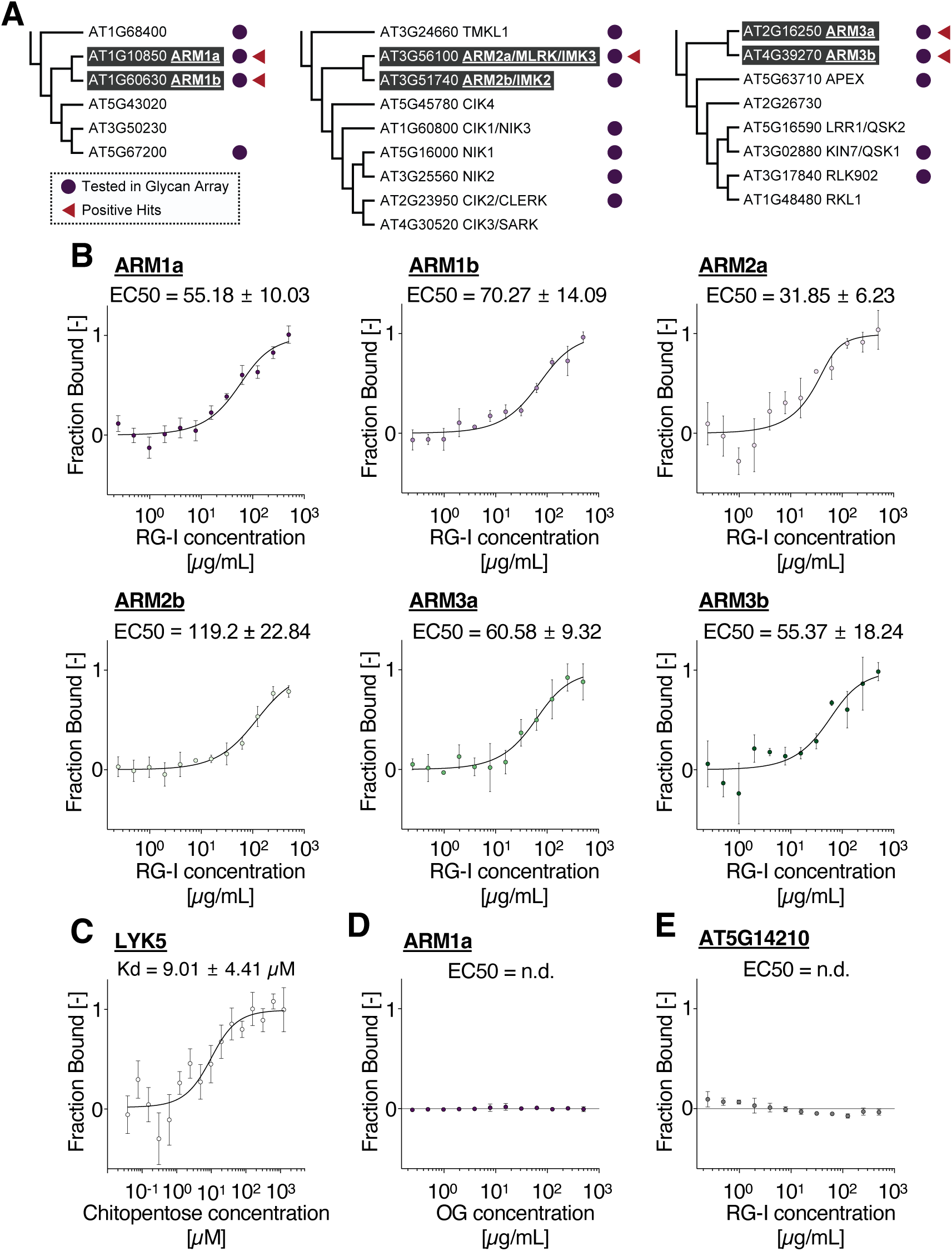
Quantification of binding between RG-I and ARM^ECD^s using Microscale thermophoresis (MST). **A.** ARM-containing clades from the receptor tree based on ECD sequences. All branches shown have bootstrap support values greater than 50. **B.** Quantification of interaction between RG-I and ARM^ECD^s using MST. Data points in the fitted-dose response binding curve indicate the fraction of labeled ARM^ECD^ bound to RG-I (mean ± SE; n=4). **C.** MST result for the interaction of chitopentaose (GlcNAc)_5_ with LYK5^ECD^ (mean ± SE; n=4). **D.** MST result for the interaction of OGs (OG^DP10-15^) with ARM1^ECD^ (mean ± SE; n=3). **E.** MST result for the interaction between RG-I and AT5G14210^ECD^ (mean ± SE; n=3).

Next, we investigated the potential PTI responses triggered by RG-I treatment. Unlike OG^DP10-15^ treatment, RG-I treatment did not induce acute Ca^2+^ influx (Fig. 3A) or production of ROIs within 120 minutes (Fig. 3B). However, RG-I did induce MAPK activation, similar to that observed with OG^DP10-15^ treatment (Fig. 3C), whereas Mock treatment did not induce MAPK activation (Supplemental Fig. 7A). Polygalacturonan (PGA) treatment, the polysaccharide form of OG^DP10-15^, resulted in weaker MAPK activation compared to RG-I and OG^DP10-15^, while the flagellin fragment flg22, also used as a positive control(Ryu et al., 2025), induced strong MAPK activation in wild-type plants but not in the flg22 receptor mutant *fls2* (Fig. 3C). Furthermore, RG-I induced stronger MAPK activation compared to GALn and ARAn (Fig. 3D), indicating that the RG-I backbone structure is critical not only for binding to ARMs but also for activating PTI responses. Root growth inhibition was observed in seedlings grown on RG-I-containing media, while no such inhibition was observed in media containing OG^DP10-15^ (Supplemental Fig. 7B and C). To explore whether these indications of immune responses are accompanied by gene expression-defined programs, we performed RNA-seq analysis to characterize genome-wide transcriptional changes triggered by RG-I treatment. These changes were compared to those induced by OG^DP10-15^ treatment (Fig. 3E and F) (Bjornson *et al*., 2021). The repertoire of differentially expressed genes (DEGs) induced by RG-I treatment was similar to that induced by OG^DP10-15^ (Fig. 3E), indicating that RG-I treatment induced a general immunity-related gene expression. In particular, the upregulated DEGs following RG-I treatment showed a more pronounced overlap with those upregulated by OG^DP10-15^ treatment compared to the overlap observed among downregulated DEGs. Among the 1,483 upregulated DEGs identified after RG-I treatment (Supplemental Fig. 8A and B), 948 genes (63.9%) overlapped with upregulated DEGs from OG^DP10-15^ treatment across the four analyzed time points, i. e. 30 and 90 minutes for both RG-I and OG^DP10-15^ treatments (Fig. 3F, upper panel). Gene Ontology (GO) analyses of the 303 upregulated DEGs shared between RG-I and OG^DP10-15^ treatment highlighted biological processes related to plant immune responses and defense mechanisms (Fig. 3G and Supplemental Table 3). Similar GO enrichment patterns for biological processes related to plant immune responses were observed for DEGs consistently upregulated at both 30 and 90 minutes following RG-I (488 genes) and OG^DP10-15^ (939 genes) treatments (Supplemental Fig. 8C and Supplemental Table 3). These transcriptomic changes were confirmed by quantitative RT-PCR analysis of the selected PTI marker gene *FLG22-INDUCED RECEPTOR KINASE 1* (*FRK1*) (Supplemental Fig. 8D). Collectively, our results suggest that RG-I treatment induces a range of PTI responses, including MAPK activation and extensive defense-related transcriptomic changes, similar to those triggered by OG^DP10-15^, but with distinct kinetics in Ca^2+^ influx and lacking ROI production.

**Figure 3.**
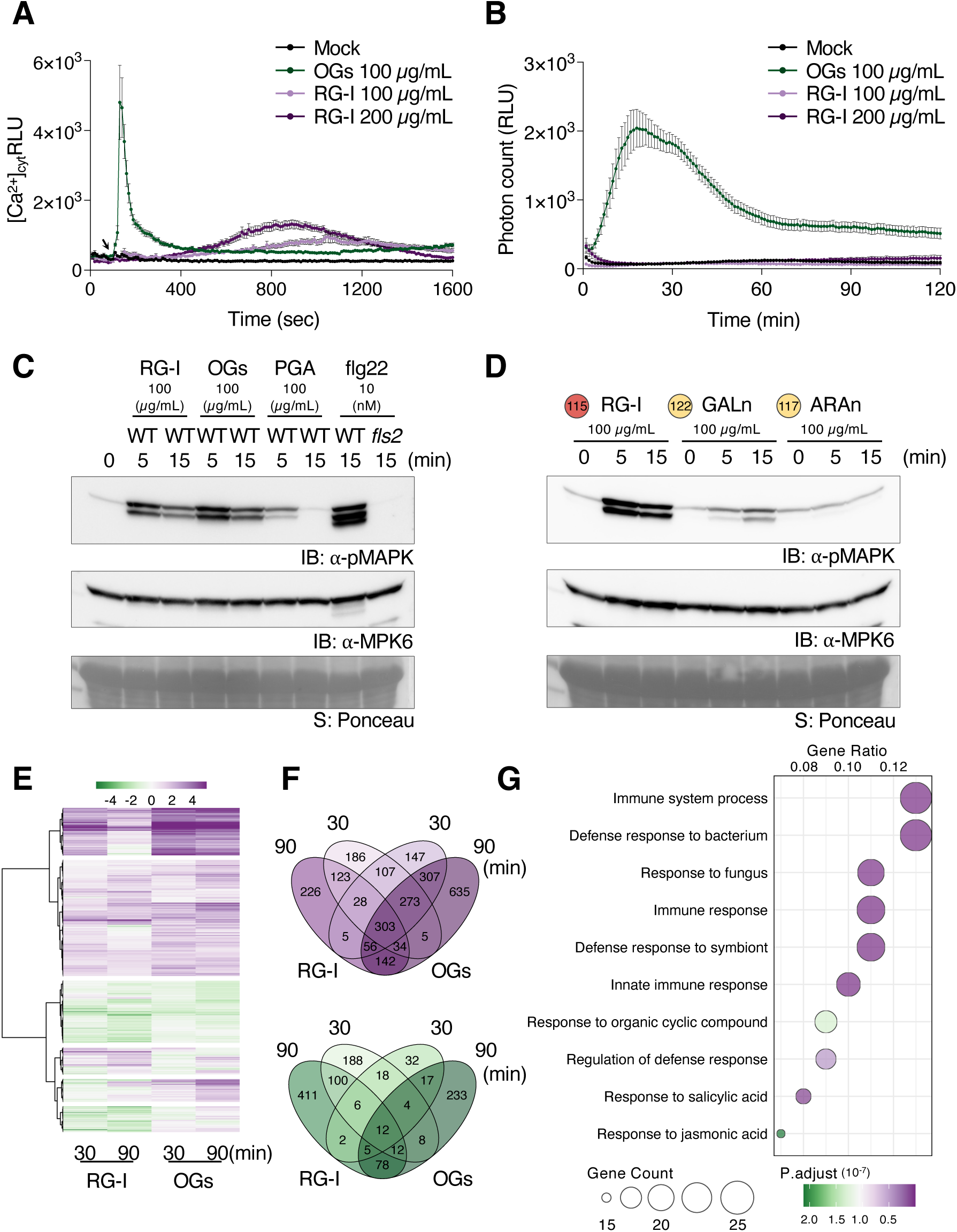
RG-I treatment induces canonical immune response outputs except for acute accumulation of cytosolic Ca^2+^ and ROI production. **A.** Cytosolic Ca^2+^ ion influx patterns were measured using the WT^AEQ^ reporter line upon treatment with either Mock, OGs (OG^DP10-15^), or RG-I (mean ± SE; n=20). The arrow indicates the treatment points (60 sec). **B.** Reactive oxygen intermediates (ROI) production kinetics after treatment with Mock, OGs, or RG-I (mean ± SE; n=24). **C and D.** Immunoblotting (IB) analysis of MAPK phosphorylation in wild-type (WT) or *fls2* mutant Arabidopsis seedlings treated with OGs, polygalacturonic acid (PGA), RG-I, or flg22 (C). IB analysis of MAPK phosphorylation in wild-type (WT) Arabidopsis seedlings treated with RG-I, Galactan (GALn) or Arabinan (ARAn) (D). Treatments, concentrations, and durations are shown on top. This experiment was repeated twice with similar results. Equal loading of protein samples on the blot was controlled by anti-MPK6 blotting (middle) and by Ponceau stain (bottom) visualizing Rubisco large subunit (rbcL). **E.** Heatmap of differentially expressed genes (DEGs) in WT seedlings treated for 30 min or 90 min with 100 µg/mL of RG-I or OGs. DEGs of OGs were retrieved from a previous study (Bjornson *et al*., 2021). **F.** Venn diagrams showing the overlap of RG-I (left) and OGs (right) DEGs. Upregulated and downregulated DEGs are shown at the top and bottom, respectively. DEG numbers are indicated in the circles. **G.** Gene Ontology (GO) analysis showing enrichment in biological process categories from overlapped 303 upregulated DEGs at both 30 min and 90 min following RG-I and OGs treatment. Bubble chart indicates the calculated gene ratio in each GO term with adjusted P-values.

To identify the essential structural requirements of RG-I for PTI activation, we performed sequential enzymatic digestion of RG-I polysaccharides to obtain RG-I oligosaccharides (Fig. 4A). First, we trimmed the side chains using galactanase and arabinofuranosidase, then removed the hydrolysis products by dialysis. The trimmed RG-I polysaccharides (RG-I Trim) still induced MAPK activation, supporting our assertion that the RG-I backbone is crucial for PTI activation (Fig. 4B). Next, we hydrolyzed the RG-I backbone using RG-I hydrolase (RG-I Digest) and purified the resulting oligosaccharides to remove proteins and unhydrolyzed polysaccharide (RG-I Oligomix). Both RG-I Digest and RG-I Oligomix induced MAPK activation, demonstrating that the activation was not due to proteinaceous molecules but rather was triggered by the RG-I oligosaccharides themselves. Additionally, when we repeated this procedure with an RG-I lyase, we found that RG-I oligosaccharides generated by either RG-I hydrolase (Hydrolase Oligomix) or RG-I lyase (Lyase Oligomix) induced MAPK activation (Fig. 4C). The hydrolase-generated oligosaccharides were then separated by size and purified by High-performance liquid chromatography (HPLC) using a PGC column to obtain defined oligosaccharides (Supplemental Fig. 9A). We successfully purified specific RG-I oligosaccharides with one or two galactose substitutions on a tetrasaccharide backbone (4+1, 4+2) and two or three galactose substitutions on a hexasaccharide backbone (6+2, 6+3) in sufficient yields for further analysis. The compositions of these oligosaccharides were determined by Matrix-assisted laser desorption ionization-time of flight mass spectrometry (MALDI-TOF-MS) (Supplemental Fig. 9B and C). Nuclear magnetic resonance (NMR) spectroscopy analysis of the 4+2 and 6+2 oligosaccharides revealed that for these two oligosaccharides each rhamnose residue was substituted with no more than one galactose (Fig. 4E and Supplemental Fig. 9D). Each purified RG-I oligosaccharide induced MAPK activation (Supplemental Fig. 9E), with the 6+2 oligosaccharide eliciting the strongest response (Fig. 4D). Next, we performed MST analyses to determine the Kd for interactions between the ARM^ECD^s and either purified oligosaccharides (4+2, 6+2) or a chemically synthesized unsubstituted RG-I backbone heptasaccharide (7+0) (Fig. 4E-H) (Romanò et al., 2024). MST analysis revealed that ARM1a^ECD^ showed increasing affinity to oligosaccharides with longer backbones and/or fewer substitutions, based on the calculated Kd values of 4+2 (162.62±135.74 µM), 6+2 (90.18±50.68 µM), and 7+0 (61.26±32.42 µM) oligosaccharides (Fig. 4F). However, the opposite trend was observed for ARM2a^ECD^ and ARM3a^ECD^, which showed decreased affinity for the 7+0 oligosaccharide compared to the 4+2 oligosaccharide (Fig. 4G and H). Collectively, our results indicate that specific RG-I oligosaccharides can function as immunogenic molecules capable of inducing MAPK activation. Furthermore, our binding studies reveal distinct oligosaccharide preferences among ARM^ECD^s, while the functional significance of these differential binding patterns requires further investigation. To investigate the loss-of-function phenotypes of ARMs, we first analyzed single mutants of each ARM gene using either transfer DNA (T-DNA) insertion lines or mutants generated by CRISPR/Cas9 technique (Supplemental Fig. 10). Additionally, we generated high-order mutants including double mutants (*arm1a/1b*, *arm2a/2b*, and *arm3a/3b*) and quadruple mutants (*arm1a/1b/2a/2b, arm1a/1b/3a/3b,* and *arm2a/2b/3a/3b*). However, none of the single, double, or quadruple mutants showed significantly different response to RG-I treatment compared to wild-type, as monitored by both MAPK activation and *FRK1* gene induction, suggesting potential functional redundancy among the ARM proteins in RG-I recognition (Supplemental Figs. 11-14). We further extended our analysis by generating and testing two different sextuple mutants (labeled #13 and #17) lacking all six ARM genes (Fig. 5A). In response to RG-I treatment, the sextuple mutants showed wild-type levels of both MAPK activation and *FRK1* expression (Fig. 5B and Supplemental Fig. 14C). The mutants also maintained normal MAPK activation when treated with RG-I oligomix or the 4+2 oligosaccharide (Fig. 5C and D). This suggests the existence of additional receptors with redundant functions, which were not tested in our initial screening or were not detected due to weak interactions below the sensitivity limit of our screening system. To further investigate this possibility, we used MST to assess interactions between RG-I and three ARM1-proximal LRR-RKs in the ECD-based receptor tree: two that were not included in our initial screening (AT5G43020 and AT3G50230) and one that was included (AT5G67200) (Fig. 5E and Supplemental Fig. 15A). MST analysis revealed that the interaction between AT5G43020^ECD^ and RG-I occurred with an EC50 value of 108.20 ± 22.68 µg/mL, which is slightly lower than that of ARM2b^ECD^, while we were unable to obtain reliable interaction values for RG-I with ECDs of two other close homologs of ARM1, AT3G50230 and AT5G67200 (Fig. 5F). Thus, we designated AT5G43020 as ARM1c. These results suggest that functional redundancy in RG-I recognition may extend beyond the six ARMs we analyzed. We also observed that RG-I induced MAPK activation was not altered in well-studied co-receptor mutants in plant immunity such as *bak1* and *cerk1*, indicating that RG-I signaling is not mediated by BAK1 or CERK1 (Supplemental Fig. 15B and C).

**Figure 4.**
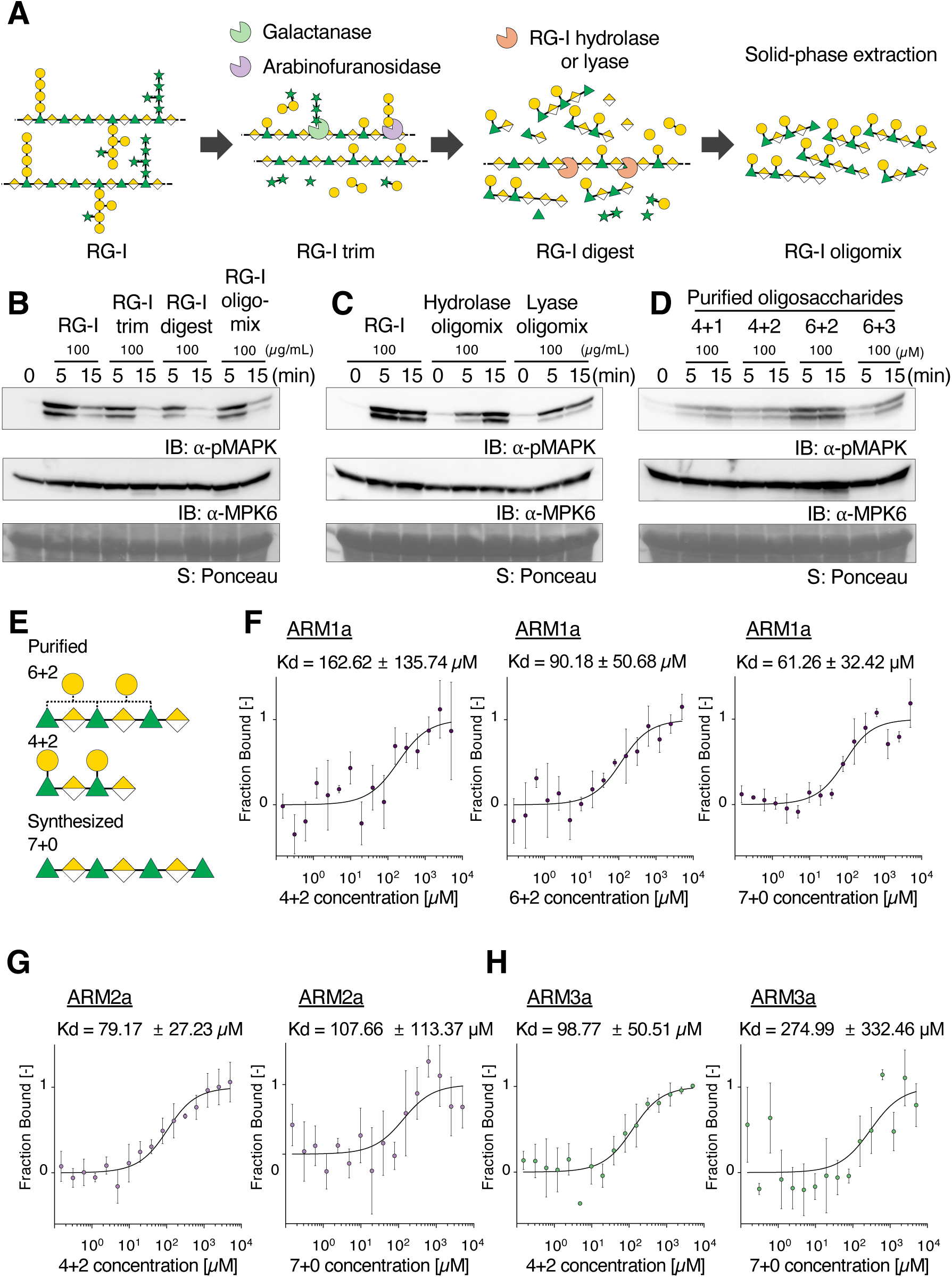
Specific RG-I derived oligosaccharides induce MAPK activation and interact with ARM^ECD^s. **A.** Schematic representation of the enzymatic digestion and preparation of RG-I fragments. **B-D.** Immunoblotting (IB) analyses of MAPK phosphorylation in wild-type (WT) Arabidopsis seedlings treated with either RG-I or RG-I fragments. Treatment conditions, including concentrations and durations, are indicated at the top of the blots. This experiment was repeated twice with similar results. Equal loading of protein samples on the blot was controlled by anti-MPK blotting (middle) and Ponceau staining (bottom), which visualizes the Rubisco large subunit (rbcL). **E.** Structures of oligosaccharides used in MST assay. **F-H**. Quantification of interaction between RG-I oligosaccharides and ARM^ECD^s using MST. Data points in the fitted-dose response binding curves indicate the fraction of labeled ARM^ECD^ bound to each oligosaccharide as described (mean ± SE; n=4).

**Figure 5.**
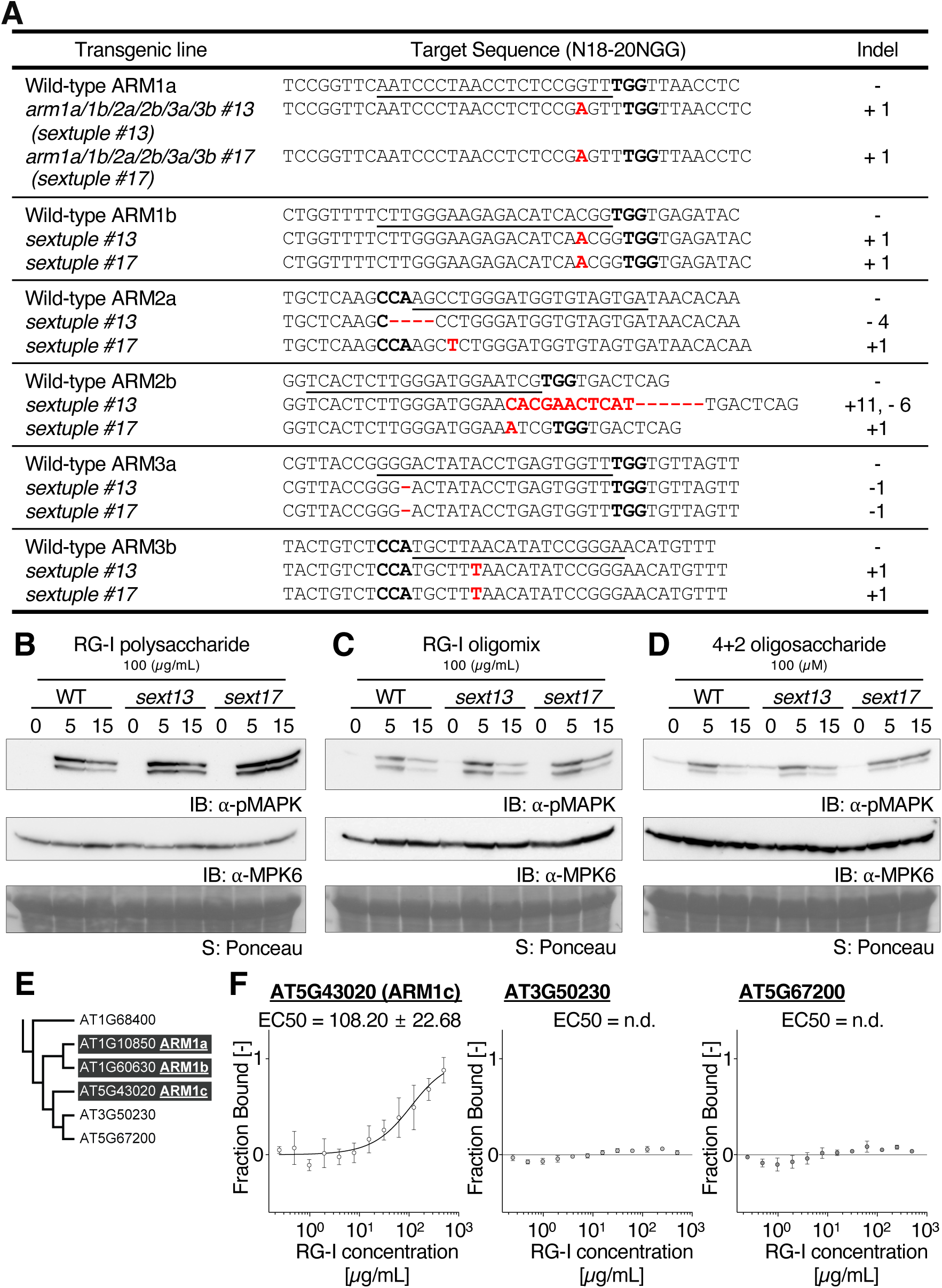
ARM sextuple mutants showed similar MAPK activation in response to RG-I compared to wild-type and possible functional redundancies with other receptors in RG-I-mediated immune responses. **A.** Gene-editing mutations identified in the six *ARM* genes corresponding to the *arm1a/1b/2a/2b/3a/3b* sextuple mutants. **B-D.** Immunoblotting (IB) analyses of MAPK phosphorylation in wild-type (WT) and *arm* sextuple mutants Arabidopsis seedlings treated with either RG-I or RG-I fragments. Treatment conditions, including concentrations and durations, are indicated at the top of the blots. This experiment was repeated twice with similar results. Equal loading of protein samples on the blot was controlled by anti-MPK blotting (middle) and Ponceau staining (bottom), which visualizes the Rubisco large subunit (rbcL). **E.** ARM1-containing clades from the receptor tree. All branches shown have bootstrap support values greater than 50. **F.** Quantification of interaction between RG-I polysaccharides and either ECDs of AT5G43020, AT3G50230, or AT5G67200 using MST. Data points in the fitted-dose response binding curves indicate the fraction of labeled ARM^ECD^ bound to each oligosaccharide as described (mean ± SE; AT5G43020, n=3; AT3G50230 and AT5G67200, n=4).

Although the sextuple mutants showed similar immune responses to RG-I treatment with regards to MAPK activation and defense-related gene induction, we found that the sextuple mutants were more susceptible to infection by the foliar pathogen *Pseudomonas syringae* pv. *tomato* DC3000 compared to wild-type plants (Fig. 6A-C). In addition, pretreatment with RG-I polysaccharide prior to infection enhanced resistance to *P. syringae* in wild-type plants, similar to the priming effect observed with flg22 pretreatment (Fig. 6D and E). Notably, only RG-I pretreatment conferred enhanced resistance in the *efr/fls2* double mutants, indicating that RG-I can function as a priming elicitor of plant defense responses, consistent with the behavior of DAMP molecules. Collectively, these findings support a role for ARMs in plant immunity, potentially through complex and redundant mechanisms involving RG-I recognition and immune priming.

**Figure 6.**
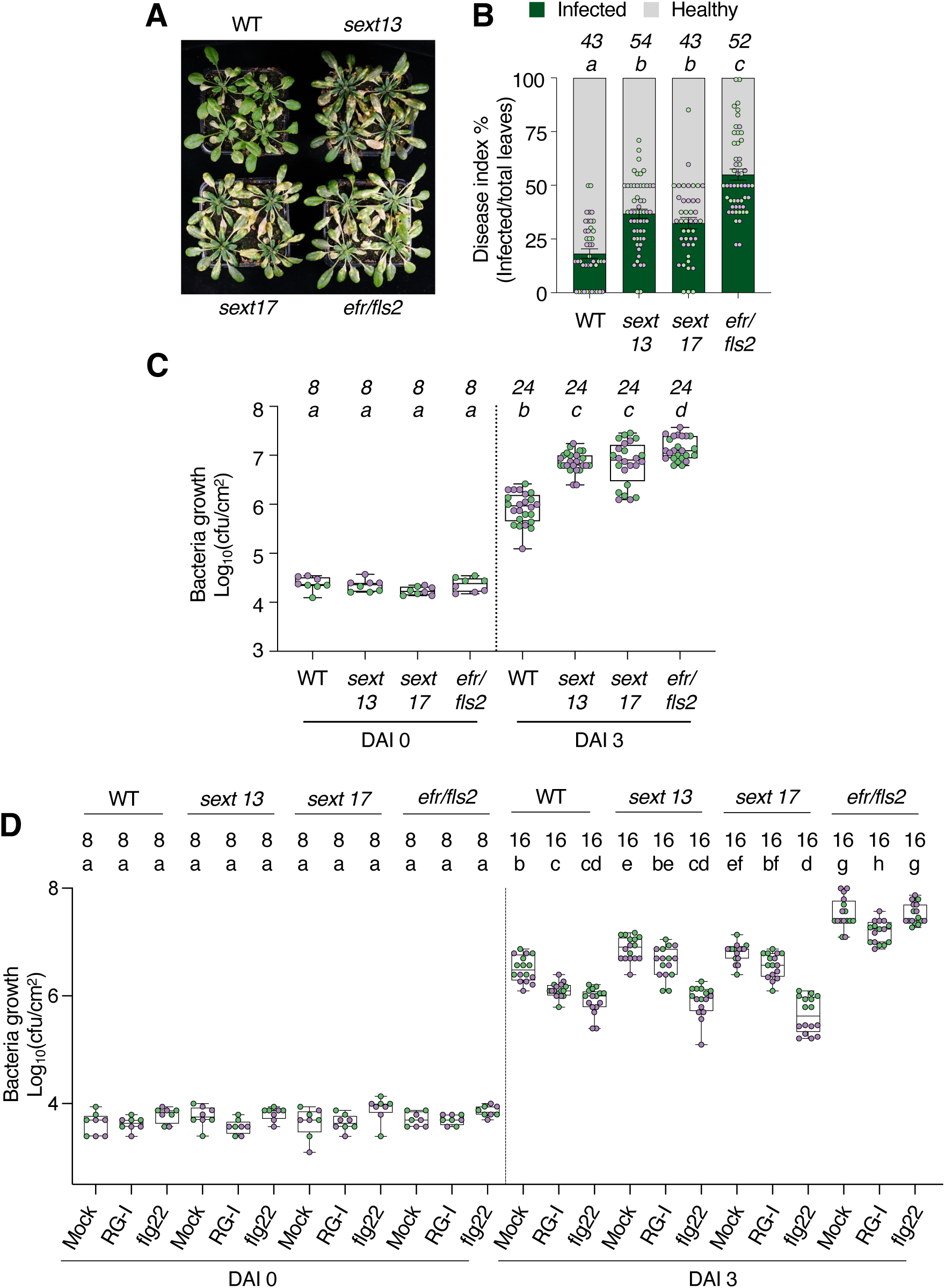
ARM sextuple mutants showed a more susceptible phenotype against *Pseudomonas syringae pv. tomato* DC3000 infection than wild-type. A. Representative pictures of Arabidopsis plants taken 5 days after inoculation with *Ps* DC3000. Genotypes are indicated at the top and bottom. **B.** The disease index percentage was calculated by dividing the number of infected leaves (more than half of the leaf being yellow) by the total number of leaves. The percentage of infected leaves is indicated with green, and that of healthy leaves is indicated with gray. Error bars indicate SE. **C.** Growth of *Ps* DC3000 in wild-type, *sextuple 13*, *sextuple 17*, and *efr/fls2.* The number of bacteria per area of leaf (cfu/cm^2^) is plotted on a log_10_ scale for day after inoculation (DAI) 0 and DAI 3. The box plot represents the first and third quartiles, centered on the median, and whiskers extend to the minimum and maximum data points. **D.** Same as (C) but growth of *Ps* DC3000 in wild-type and *efr/fls2* leaves pretreated with Mock, RG-I (100 µg/mL), or flg22 (100 nM) one day prior to bacterial inoculation. The box plot represents the first and third quartiles, centered on the median, and whiskers extend to the minimum and maximum data points. **B-D.** Numbers of biologically independent observations (*n*) are indicated above the letters, and dots with different colors (green and purple dots) indicate different experimental trials. Letters above the boxes (a-d) indicate the results of statistical analysis using a linear mixed effect model with one-way ANOVA followed by a Tukey’s multiple comparison test (two-sided, adjusted for multiple comparisons, *p* < 0.05).

## Discussion

In land plants, pectin is a major component of the primary cell wall, accounting for approximately 30% of its dry weight (Willats et al., 2001). Pectin consists of a group of highly complex and heterogeneous cell wall polysaccharides, organized into five major domains: homogalacturonan (HG), RG-I, RG-II, apiogalacturonan, and xylogalacturonan (Harholt et al., 2010). While the exact composition varies with environment, tissue type, and plant species, HG typically makes up around 60% of pectin and RG-I contributes 20% to 30% (Mohnen, 2008). Although HG and RG-II are known to be involved in strengthening the cell wall, the specific function of RG-I remains less understood (Harholt *et al*., 2010). In this study, we identified a new role of RG-I in plant defense responses, similar to that of OGs from HG or cellooligomers from cellulose, which function as DAMPs. RG-I specifically activates MAPK and induces defense genes without triggering acute Ca^2+^ influx or ROI production, unlike OGs and cellooligomers, which trigger all these responses (Fig. 3). These differences suggest that RG-I may engage a distinct set of downstream signaling components, or that the RG-I oligosaccharides used in this study may lack key structural features necessary for the full spectrum of ARM receptor activation. However, unlike OGs, RG-I induces slight root growth inhibition, which may reflect differential receptor expression patterns between ARMs and WAKs, or differences in the signaling modules they mediate to balance growth-defense trade-offs. Taken together, the distinct responses may also represent a temporal sequence of pectin degradation, where less structurally complex HG is degraded earlier to release OGs that trigger acute responses, while the more complex RG-I produces oligosaccharides later to maintain prolonged defense signaling and reinforce growth-defense trade-offs during extended pathogen pressure. Furthermore, in natural contexts, oligosaccharides released from pectin components could vary depending on cell wall composition in a tissue-specific manner or on the diverse arrays of cell wall-digesting enzymes produced by different pathogens. Comprehensive profiling of tissue- and pathogen-specific oligosaccharide release patterns will provide crucial insights into the physiological relevance of these diverse cell wall-derived DAMP recognition systems.

We identified that oligosaccharide fragments of RG-I can bind to ARM^ECD^s and induce MAPK activation in Arabidopsis (Fig. 4). However, these oligosaccharides were less effective than RG-I polysaccharides (Supplemental Fig. 9E), which may reflect suboptimal structural features of tested RG-I oligosaccharides. Given that longer OGs (DP 10-15) elicit stronger immune responses than shorter OGs, the RG-I fragments we tested could be less effective due to their shorter length and limited side-chain substitutions. We hypothesize that longer RG-I oligosaccharides with more defined substitution patterns may be required to achieve optimal ARM receptor activation and the full spectrum of immune responses. While both RG-I lyase and RG-I hydrolase produced RG-I oligosaccharides that activated MAPK similarly (Fig. 4C), further research is needed to understand which oligosaccharides are predominantly produced during biotic interactions in a natural context. Similarly to the profiling of OG fragments in Arabidopsis upon *Botrytis cinerea* infection (Voxeur et al., 2019), profiling RG-I fragments from interactions with microorganisms that contain RG-I lyases in their genomes, such as *Aspergillus niger* or *Bacillus subtilis* (Khosravi et al., 2017; Ochiai et al., 2007), needs to be analyzed in future studies. Additionally, during the enzymatic digestion to fractionate RG-I oligosaccharides, we observed that in most oligosaccharides, at least one rhamnose residue is substituted with a single galactose. This suggests that galactose-substituted oligosaccharides might also be generated in a natural context, indicating potential functional significance. Chemically synthesizing these RG-I oligosaccharides will allow us to investigate their specific function (Romanò *et al*., 2024).

The conventional approach to exploring novel ligand-receptor pairs involves first identifying the immunity-eliciting molecule and then finding its cognate receptor through forward genetics screening. However, this method faces significant limitations, particularly due to the functional redundancies of PRRs, except in rare cases where dominant negative mutations are generated. Our high-throughput screening pipeline was developed to overcome these limitations. Through the pipeline reported here, we identified five ARM receptors that interact with RG-I and its fragments, and additionally found that AT3G51740 (ARM2b) and AT5G43020 (ARM1c) showed reduced interaction activity with RG-I in MST analysis (Figs. 1D, 2B, and 5F). However, our screening approach also revealed inherent limitations of high-throughput glycan array-based interaction systems. While WAK1 is known to interact with PGA and OGs, we did not observe such interactions in our array. This is likely due to the short length and methylesterification of the printed OGs^DP6^ and ineffective immobilization of PGA on the array. Moreover, the relatively low hit rate observed in our study likely reflects multiple technical and biological factors. First, using unnormalized secreted ECD proteins from crude insect cell supernatants leads to variable protein concentrations across the library, with some ECDs potentially present at concentrations insufficient for reliable detection. Additionally, the presence of other secreted proteins and metabolites in the crude supernatants could interfere with specific binding interactions. Second, the detection sensitivity of the array may not have been sufficient to capture weak interactions; for instance, ARM2b showed no array signal but bound RG-I in MST assays. Third, immobilizing glycans via their reducing ends could mask interactions that depend on this terminus. Finally, some receptors may require regions beyond the annotated ECD for proper folding or ligand engagement. While these limitations resulted in a conservative hit rate, our approach successfully prioritized the most robust receptor-ligand interactions for detailed validation.

Notably, while LRR domains have traditionally been associated with proteinaceous ligand binding in plants, Toll-like receptors (TLRs) in animals, which also contain LRR domains, recognize a wide range of ligands, including lipopolysaccharides, peptidoglycans, and nucleic acids (Kawai et al., 2024). Similarly, recent studies in plants suggest that LRR-containing receptors also engage with non-proteinaceous ligands, such as CORK1/IGP1-4 receptors with cello-oligomer ligands (Martín-Dacal *et al*., 2022; Tseng *et al*., 2022). In addition, LRX8, an LRR extensin protein, can interact with de-esterified OGs in the presence of Rapid Alkalization Factor 4 (RALF4) peptides (Moussu et al., 2023). Together with these examples, our study supports the notion that plant LRR domains may have broader ligand recognition capabilities than previously assumed (Lee et al., 2024).

Although the *arm* sextuple mutants still exhibited MAPK activation and *FRK1* gene induction upon RG-I treatment (Fig. 5 and Supplemental Fig. 14C), these mutants were less resistant to pathogen infection (Fig. 6). While this increased susceptibility to pathogen infection could result from indirect effects, such as altered plant fitness and metabolism in the high-order mutations, our findings are consistent with a possible role for ARM receptors in plant immunity. Notably, the retention of MAPK activation and *FRK1* gene induction in the sextuple mutants is reminiscent of recent findings on WAK receptors upon OGs perception. Indeed, analysis of *wak1-5* quintuple knock-out mutants suggests that additional receptors besides WAKs, including WAK-like (WAKL) family RKs, might be involved in the OGs response (Herold *et al*., 2024). Similarly, we propose that additional redundant ARM receptors exist that were not tested in the screening due to either protein library limitations or insufficient affinity for glycan array detection. For example, ARM1c was not included in the initial screening but, when tested later, showed RG-I binding affinity (Fig. 5F). Therefore, we hypothesize that the maintained MAPK induction in the sextuple mutant might result from additional redundant ARM receptors that participate in Arabidopsis RG-I signaling.

ARM receptors have distinct tissue-specific and developmental stage-specific expression patterns (Supplemental Fig. 16) (Mergner et al., 2020; Wu et al., 2016). These patterns may indicate a sophisticated regulatory system, potentially enabling ARM receptors to monitor cell wall dynamics in a spatially and temporally coordinated manner. Given that pectin dynamics are tightly regulated for cell wall integrity maintenance and cell morphogenesis (Du et al., 2022), this diversity in expression likely enables robust surveillance of cell wall status across different tissues. This spatially coordinated expression of multiple receptors is reminiscent of the high-order regulation observed in the CLAVATA (CLV) signaling pathway, where BAM receptors exhibit tightly controlled spatial expression patterns regulated by CLV3-CLV1, leading to precise feedback control of stem cell populations (Nimchuk et al., 2015). Similarly, ARM receptor redundancy might reflect a system optimized for tissue-specific and context-dependent signaling, allowing plants to adapt to diverse physiological and environmental conditions. Together with the WAK-mediated OG recognition mechanisms, our findings uncover an additional layer of surveillance through ARM receptor-mediated RG-I recognition, which collectively establishes a sophisticated system for monitoring cell wall integrity via distinct pectin-derived DAMPs.

Our previous application of the high-throughput microarray approaches demonstrated their versatility in detecting ligand-receptor interactions by diversifying the ligand range for individual plant receptors (Lee et al., 2024; Parys et al., 2021). Here, we expanded this approach by screening an extended plant receptor library against a collection of glycan molecules, including oligo- and polysaccharides. Our results validated this high-throughput screening pipeline for identifying novel ligand-receptor pairs. This versatile screening pipeline could be further expanded to explore crop plant receptor libraries and targeted glycan ligand libraries from specific structures of cell walls in fungi, oomycetes, or parasitic plants, paving the way for translational research.

## Methods

### Plant materials and growth conditions

*Arabidopsis thaliana* accession Columbia (Col-0) was used as the wild-type and used for transgenic line generation. The mutant lines *cerk1* (SALK_007193), *bak1-4, bak1-5,* and *efr/fls2* were previously described (Miya *et al*., 2007; Schwessinger et al., 2011; Smakowska-Luzan *et al*., 2018). The single T-DNA mutant lines *arm1a-2* (GABI_210H07), *arm1b-2* (GABI_510G08)*, arm2a-1* (SALK_024031)*, arm2a-2* (SAIL_600F09)*, arm2b-2* (SALK_029864)*, arm3a-1* (SALK_011390)*, arm3a-2* (SALK_025297C)*, arm3b-1* (SALK_071306), and *arm3b-2* (SALK_072419C) were genotyped using PCR and checked for full-length transcripts using RT-PCR, both with corresponding primers (Supplemental Table 4). Arabidopsis seeds were surface-sterilized and grown on Petri dishes containing ½ Murashige and Skoog (MS) medium with Gamborg B5 vitamins (Duchefa, M0231), 1% sucrose (Roth), and 0.8% agar (AppliChem), adjusted to pH 5.7 with KOH, in a growth room at 22 °C under long-day light conditions (16 h light/8 h dark). For the pMAPK, qPCR, or RNAseq experiments, 5-day-old Arabidopsis seedlings vertically grown on plates were transferred to a 6-well plate containing 2 mL of liquid media (½ MS including MES buffer (Duchefa, M0255) and 1% sucrose) and incubated further for 1 week. For the infection assays, Arabidopsis seeds were grown on soil in a growth chamber at 22 °C under day-neutral light conditions (12 h light/ 12 h dark).

### Receptor tree

Protein sequences for receptors were obtained from a previous publication (Ngou et al., 2022), while additional receptor proteins were obtained from TAIR (TAIR10_pep_20101214). A total of 518 candidate receptors were selected based on the functional domains identified by Pfam database [PMID: 39540428]. Transmembrane domains and transit peptides were analyzed using DeepTMHMM (Hallgren et al., 2022). Then, receptor candidates without transmembrane domains or with an ECD length of less than 50 amino acids were excluded. The full protein sequences of 488 receptors were aligned with MAFFT (Rozewicki et al., 2019). Maximum likelihood trees were inferred with IQ-Tree, and support values for individual nodes were estimated using 1,000 pseudoreplicates of the ultrafast bootstrap approximation(Minh et al., 2020). Trees with additional labels were visualized by iTOL v6 (Letunic and Bork, 2024).

### S2 cell expression

Recombinant ECDs of LRR-RK in pECIA2 expression vector, that provides a C-terminal Fc-tag, V5-tag and His-tag, were described previously (Smakowska-Luzan *et al*., 2018), and additional ECDs were cloned into pECIA2 in this study. ECD expression was performed as previously described (Clasen et al., 2023). Briefly, expression clones were transiently transfected into *Drosophila* S2 cells (ExpreS2ion Biotechnologies) using the Expres2 TR transfection reagent (ExpreS2ion Biotechnologies) and cultured at 27 °C. Upon transfection, the culturing temperature was switched to 25 °C. After 24 hours from transfection, protein expression was induced with 1 mM CuSO_4_ for a further 72 hours. After expression, the supernatant including the secreted ECDs was collected and supplemented with cOmplete™, EDTA-free Protease Inhibitor (Roche, 11873580001) and 0.02% NaN_3_. Protein expression was determined by immunoblotting using anti-V5-HRP antibodies (Invitrogen, R962-25).

### Glycan array

The plant glycan microarray was printed as previously described (Ruprecht *et al*., 2020). In brief, the synthetic oligosaccharides and commercially available polysaccharides were dissolved in the coupling buffer (150 mM sodium phosphate, pH 8.5) in a concentration of 200 µM for oligosaccharides and 200 µg/mL for polysaccharides (chitosan and 1,3-linked glucan at 20 µg/mL due to low solubility), and printed in triplicates (each drop was about 0.5 nl) on CodeLink N-hydroxyl succinimide (NHS) ester-activated glass slides (SurModics Inc., USA) using a non-contact piezoelectric spotting device (S3; Scienion, Germany). Sixteen identical fields as depicted in Supplemental Fig. 3 with all glycans in triplicates were printed on a glass slide at RT. After printing, the remaining NHS groups on the microarray slides were quenched for 1 hour at RT by immersing them in 100 mM ethanolamine, 50 mM sodium phosphate, pH 9. Then, the slides were washed three times by immersion in deionized water, dried and stored at 4 °C until usage. For the ECD binding assay, a 16-well grid was applied to the glass slide and the array was blocked for 45 min with 200 µL 1 % bovine serum albumin (BSA) in binding buffer (20 mM HEPES, 100 mM NaCl, 1 mM CaCl_2_, 1 mM MnCl_2_, 1 mM MgCl_2_, pH 6.8) per well. Next, the 100 µL of the supernatant of the insect cell media including the expressed ECDs was applied and incubated for 1 hour. After washing three times for 5 min with 200 µL binding buffer, 100 µL of a solution with a polyclonal goat anti-human IgG antibody labeled with Alexa Fluor 647 (Invitrogen, A21445) diluted 1:400 in 1 % BSA in binding buffer was applied for 45 min. Then, the unbound secondary antibody was removed by washing once for 5 min with 200 µL binding buffer including 0.1% Tween20, once 5 min with binding buffer, and rinsing the slide twice with water. All steps were performed at RT. The slides were scanned on a Genepix 4300A microarray scanner (Molecular Devices) using the 635 nm laser at photomultiplier gain PMT800 for initial screening and additionally PMT500 for the further analyzing six ARM proteins. The images were obtained and analyzed using the GenePix Pro 7 software.

### On-array enzymatic digestion

For on-array enzymatic digestion, RG-I hydrolase (0.13 mg/mL in 100 mM sodium acetate buffer, pH 5.5), arabinofuranosidase, or galactanase (10 U/mL in 100 mM sodium acetate buffer, pH4) were incubated in 100 µL reactions per well for 16 hours at 40°C. After washing twice with binding buffer, antibody staining using INRA-RU2 was performed as previously described (Ruprecht *et al*., 2017) and the ARM1a^ECD^ binding was conducted as described above.

### Polysaccharides and oligosaccharides

The sources of the purchased polysaccharides are indicated in Supplemental Table 2. The chemical synthesis of the oligosaccharides was previously described (Bartetzko et al., 2015; Bartetzko et al., 2017; Dallabernardina et al., 2016; 2017; Schmidt et al., 2015; Senf et al., 2017; Zakharova et al., 2013). The branched RG-I oligosaccharides were prepared as follows: The side chains of the RG-I polysaccharide (5 mg/mL, Megazyme) were trimmed using arabinofuranosidase (AFASE, Megazyme, 5 U/mL) and galactanase (EGALN, Megazyme, 5 U/mL) in 100 mM sodium acetate buffer pH 4 at 40°C for 16 hours. After removing the arabinan and galactan fragments using dialysis (SnakeSkin dialysis tubes, 3.5 MWCO) for 8 hours at 4°C, exchanging the water three times, the resulting “RG-I trim” was digested using RG-I hydrolase (13 µg/mL) for 16 hours at 40°C in 100 mM sodium acetate pH 5. These “RG-I digest” still included hydrolytic enzymes that were heat-inactivated for 10 min at 85 °C. To obtain pure oligosaccharides, the sample was loaded on Carbograph solid phase extraction (SPE) columns (S*Pure) and was eluted using acetonitrile/0.25 % ammonium formiate buffer, pH 3 (1:1). Aliquots of the different samples in the purification process and the final “RG-I oligomix (Hydrolase oligomix)” were lyophilized. The same procedure was used to purify the “Lyase oligomix”, except that RG-I lyase (18 µg/mL) was used. To analyze the composition of the RG-I hydrolase oligosaccharides a Shimadzu LC-MS system was used with a Hypercarb porous graphitized carbon (PGC) column (Thermo Fisher, 4.6x100 mm, 5 µm particle size) equipped with an Alltech 3000 Evaporative Light Scattering Detector (ELSD) and a Shimadzu LCMS-2020 mass spectrometer. To obtain defined individual oligosaccharides the RG-I oligosaccharide mixture was separated using HPLC with a semipreparative Hypercarb PGC column (Hypercarb, 10x100 mm, 7 µm particle size), The flow rate was 2 mL/min and the oligosaccharides were eluted applying a water (with 0.25 % ammonium formiate, pH 3), acetonitrile gradient starting with 5% acetonitrile (ACN) for 8 minutes followed by a ramp up to 30% ACN for 60 minutes, a short increase to 100% ACN for 4 minutes, a decline back to 5% and an equilibration at 5% ACN for 5 minutes. The fractions were collected in 30 seconds intervals and analyzed with LC-MS as described above and with MALDI-TOF-MS using the Autoflex Speed device (Bruker Daltonics, Germany) operated by the FlexControl 3.4 software. Fractions with the same oligosaccharide were pooled and the structures of the purified oligosaccharides were determined using nuclear magnetic resonance (NMR) analyses.

### NMR analyses

Experiments were conducted on a Bruker III Avance 600 MHz spectrometer using Topspin 3.6.5 software. Spectra were recorded in D_2_O at 300 K and referenced externally for ^1^H to DSS in D_2_O (0 ppm) and for ^13^C to dioxane in D_2_O (67.4 ppm). Standard pulse programs and settings from the Bruker library were employed unless otherwise stated. For recording HSQC spectra, a F1 FIDRes of 62 Hz (TD 1024) was achieved using non-uniform sampling with a NUS percentage of 40%. For HMBC, a F1 FIDRes of 65 Hz (TD 1024) was achieved using non-uniform sampling with a NUS percentage of 50%. Spectra were processed and assigned using Bruker Topspin 3.6.5.

### ARM extracellular domain expression and purification

ARM1a^ECD^ (residues 31-253), ARM1b^ECD^ (24-247), ARM2a^ECD^ (53-406), ARM2b^ECD^ (39-455), ARM3a^ECD^ (32-443), ARM3b^ECD^ (29-413), AT5G14210^ECD^ (27-393), AT5G43020/ARM1c^ECD^ (29-248), AT3G50230^ECD^ (34-300), AT5G67200^ECD^ (19-281), and LYK5^ECD^ (27-276) sequences were amplified from Arabidopsis cDNA, cloned into the pB1-Mel vector (Invitrogen) with a C-terminal TEV protease site, StrepII tag, BamHI site (Gly-Ser), and 9×His tag, and used for Bacmid generation. After viral amplification, recombinant ECD proteins were expressed in *Trichoplusia ni* High Five insect cells (Expression Systems) at 21 °C for 72 hours. The secreted proteins were purified using Ni-NTA affinity chromatography (HisTrap HP, Cytiva) with binding buffer (20 mM NaH_2_PO_4_/Na_2_HPO_4_ buffer pH 7.5, 500 mM NaCl, and 20 mM Imidazole) and subsequently size-exclusion chromatography (SEC) (Superdex 200 Increase 10/300 GL, Cytiva) with SEC buffer (20 mM NaH_2_PO_4_/Na_2_HPO_4_ buffer pH 7.5 and 200 mM NaCl). Purification yields were: ARM1a^ECD^ 3.0 mg, ARM1b^ECD^ 3.6 mg, ARM2a^ECD^ 0.5 mg, ARM2b^ECD^ 1.0 mg, ARM3a^ECD^ 6.9 mg, ARM3b^ECD^ 0.3 mg, AT5G14210^ECD^ 0.4 mg, AT5G43020/ARM1c^ECD^ 0.5 mg, AT3G50230^ECD^ 4.6 mg, AT5G67200^ECD^ 1.4 mg, and LYK5^ECD^ 0.4 mg per liter of culture.

### MicroScale Thermophoresis (MST) assay

The purified ECD proteins were labeled with the Monolith His-Tag Labeling Kit RED-tris-NTA 2^nd^ Generation (MO-L018, NanoTemper) according to the manufacturer’s instructions. 50 nM of labeled proteins and varying concentrations of ligands were mixed in MST buffer (20 mM NaH_2_PO_4_/Na_2_HPO_4_ buffer pH 7.5, 200 mM NaCl, 0.005% Tween-20), and then loaded into Monolith Premium Capillaries (MO-K025, NanoTemper). Measurements were performed at 25 °C in a Monolith NT.115 instrument with LED power set to 80% and MST power to 20% or 40%. Measurements were repeated at least three times. The data were analyzed by plotting the log_10_ values of ligand concentrations against the change in normalized thermophoresis (ΔFnorm [‰]) using MO.Affinity Analysis software (NanoTemper). The data were then analyzed with GraphPad Prism 10 software to fit curves and determine the EC50 or Kd values.

### pMAPK assay

Elicitor treatment was performed by replacing the previous media in the 6-well plates with 2 mL of sterile MilliQ water containing each elicitor and incubated at 22 °C for indicated time points. 16 seedlings were used for total protein extraction. Total proteins were extracted in 300 µL of extraction buffer (50 mM Tris-Cl pH 7.5, 150 mM NaCl, 2 mM Na_2_MoO_4_, 2 mM Na_3_VO_4_, 5 mM NaF, 2 mM phenylmethylsulfonyl fluoride, 1 mM DTT, 10% Glycerol, 1% IGEPAL, cOmplete™, EDTA-free Protease Inhibitor (Roche)). 40 µL of protein extract was mixed with SDS-PAGE loading buffer, boiled at 90 °C for 2 minutes, and subjected to SDS-PAGE. Immunodetection was performed using anti-p44/p42 MAPK antibodies (9102, Cell Signaling; 1:5000), anti-MPK6 antibodies (AS122633, Agrisera; 1:5000), and anti-Rabbit-HRP antibodies (A6154, Sigma-Aldrich; 1:5000). Signals were detected with a ChemiDoc instrument (Bio-Rad).

### Reactive oxygen intermediate (ROI) burst assay

4 mm leaf discs from 6-week-old wild-type plants were placed in a 96-well luminescence assay plate, vacuum-infiltrated in sterile MilliQ water and incubated overnight. The water was replaced with a treatment solution containing elicitors, L-012 (Wako Chemicals), and horseradish peroxidase (Thermo Fisher Scientific). Relative luminescence was measured every minute for 120 min using a BiTec Synergy 4 microplate reader and summed for total photon counts.

### Ca^2+^ influx measurement

Col-0^AEQ^, carrying the calcium reporter aequorin protein, were grown in a 96-well luminescence assay plate in liquid media (½ MS and 1% sucrose, adjusted pH 5.7) for 7 days. A day before the treatment, the media was replaced with 100 µL of 10 µM coelenterazine h (Biosynth, FC36332) solution and incubate for overnight in the darkness. Before treatment, the plates were scanned for 1 min (300 msec per well with 6 sec interval) to obtain resting level readings using a Spark Cyto microplate reader (TECAN). For treatment, 50 µL of 3-fold concentrated elicitor solutions were added to the plates, and further scans were conducted for 25 min.

### RNAseq analysis

12-day-old Arabidopsis seedlings grown in 6-well plates were treated for 0min, 30 min, or 90 min with 100 µg/mL of RG-I polysaccharide. Total RNA was isolated from pools of 16 seedlings using the RNeasy Plant Mini Kit (QIAGEN). All samples were prepared in triple replication from independent experiments. Five hundred nanograms of total RNA from each sample was used for cDNA and NGS library preparations with the QuantSeq 3’ mRNA-Seq Library Prep kit (FWD) (Lexogen, 015). The individual NGS libraries were pooled in equimolar ratios and sequenced on a NovaseqS4 in single end-150 base mode. The quality check of raw reads was conducted by FastQC (ver 0.11.9). The reads were processed and quantified against the reference transcriptome (TAIR10.transcripts.fa) using Kallisto (ver 0.46.2). The Kallisto-processed reads from OGs treatment were retrieved from a previous study (Bjornson *et al*., 2021) using getDEE2 (ver 1.14.0). The abundance results were analyzed using DESeq2 (ver 1.44.0). Differentially expressed genes (DEGs) were identified with a cut-off of |Log2FC| > 1 and Padj < 0.05. These DEGs were then used to generate a heatmap with the pheatmap package (ver 1.0.12) and a Venn diagram with the ggvenn package (ver 0.1.10). Gene Ontology (GO) analysis was performed using the enrichGO function from the clusterProfiler package (ver 4.12.2) to assess biological processes (BP). Enriched functions were emphasized with a bubble-chart, excluding the hypoxia-related gene sets considered as artifacts of the treatment method to avoid misinterpretation.

### RT-qPCR analysis

Elicitor solution treatment was performed by replacing the previous media in the 6-well plates with new sterile MilliQ water containing each elicitor. After replacement, the plates were vacuum infiltrated for 10 min and incubated 1 hour or 2 hours. 8 or 16 seedlings were used for total RNA extraction. Total RNA was isolated using the GeneMATRIX Universal RNA Purification Kit (EURx). Subsequently, one microgram of isolated total RNA was used for cDNA synthesis with the RevertAid First Strand cDNA Synthesis Kit (Thermo Fisher Scientific). qPCR analyses were performed with Takyon ROX SYBR 2X MasterMix dTTP blue (Eurogentec, UF-RSMT-B0701) with target primers (Supplemental Table 4) using a Bio-Rad CFX Opus Real-Time PCR system. Actin2/8 cycle threshold (Ct) was used as an endogenous control and ΔCt of mock treatment for each genotype was used for normalization.

### CRISPR/Cas9 genome editing

CRISPR/Cas9 genome editing was performed as previously described (Richter et al., 2018). Briefly, the selected guide RNA (gRNA) sequences targeting the ARM genes were inserted into the guide RNA scaffold (sgRNA) within GreenGate entry vectors. The annealed reverse and forward complement sequences of the gRNA primers, each with different overhangs (Supplemental Table 4), were ligated into the entry vector cut with BpiI (New England Biolabs, FD1014). ARM1a sgRNA1 and ARM1b sgRNA2 entry clones were assembled into the destination vector pGGZ003 using the GreenGate reaction, together with entry clones containing the *Petroselinum crispum* ubiquitin4-2 promoter, codon-optimized Cas9, *Pisum sativum* pea3A terminator, and seedcoat-localized YFP (pAlligator:YFP). The same process was applied to ARM2a sgRNA1 and ARM2b sgRNA2, as well as to ARM3a sgRNA1 and ARM3b sgRNA2. Each destination vector was introduced into *Agrobacterium tumefaciens* (GV3101 pSoup). Wild-type Arabidopsis was then transformed with these destination vectors to generate single or double mutants using the floral dip method. YFP-positive T1 seeds were grown on soil, and PCR was performed using the corresponding primer sets (Supplemental Table 4). The PCR products were sequenced with the Sanger method and analyzed with the Tide program (http://shinyapps.datacurators.nl/tide/). YFP-negative T2 seeds were selected to obtain plants lacking the Cas9 cassette and analyzed by sequencing using the same methods. Higher-order mutants, such as quadruple or sextuple mutants, were generated by introducing additional gRNA sets into the double mutants or quadruple mutant, respectively.

### Bacterial infection assays

4-week-old Arabidopsis plants grown on soil were sprayed with a *Pseudomonas syringae pv. tomato* DC3000 inoculum at OD_600_ = 0.002 in 10 mM MgCl_2_ supplemented with 0.02% Silwet L-77. For the priming experiments, 4-week-old Arabidopsis plants grown on soil were pretreated by infiltration with mock, 100 µg/mL RG-I, or 100 nM flg22 using a 1 mL needleless syringe, and incubated for 1day. A *P. syringae* inoculum at OD_600_ = 0.0002 diluted in 10 mM MgCl_2_ was then infiltrated in the pretreated leaves. Three days after inoculation, 8 leaf discs from 4 independent plants were bored, ground, and homogenized in 10 mM MgCl_2_. The bacterial titer was assessed by serial dilution plating. Representative picture was acquired five days after inoculation. The disease index was calculated by dividing the number of infected leaves (more than half of the leaf being yellow) by total number of leaves.

### Data visualization and statistical analysis

GraphPad PRISM 10 and R (version 4.5.1 with Bioconductor) were used for data analysis and visualization. Statistical analyses were performed using either one-way ANOVA (two-sided) followed by Tukey’s multiple comparisons test in GraphPad PRISM, or linear mixed-effects modeling followed by one-way ANOVA (two-sided) with Tukey’s multiple comparisons test in R. For linear mixed-effects modeling, we used the lme4 package (version 1.1-35.5), with wild-type control or mock treatment as fixed effects and experimental replicates as random effects, using the formula: Value ∼ treatment + (1|Trial). Values were log_10_-transformed when this improved the fit to the linear model. Different letters above the plot indicate significant differences between groups based on all pairwise comparisons using multiple comparisons tests.

## Materials availability

All materials generated in this study are available from the corresponding authors upon reasonable request for non-commercial research purposes.

## Data availability

The RNA-seq datasets generated and analyzed in the current study have been deposited in the Sequence Read Archive at NCBI under bioproject accession code PRJNA1195916. All source codes that we used in this study are also available in *Zenodo* (DOI: 10.5281/zenodo.14760738).

## Author contributions

**Project Administration,** Y.B. and F.P.; **Conceptualization,** D.H.L., C.R., Y.B., and F.P.; **Investigation,** D.H.L, C.R., J.M.L, M.S.C, T.H., H.R., S.C., G.H., N.E., B.E., and M.B.; **Supervision,** D.H.L., C.R., Y.B., and F.P.; **Methodology,** D.H.L., C.R., J.M.L, M.S.C, T.H., H.R., S.C., G.H., N.E., B.E., M.B., C.H.C., H.S.L., and Y.B.; **Resources,** D.H.L., C.R., Y.B., E.S.L., and F.P.; **Software,** D.H.L., C.R., and F.P.; **Formal Analysis,** D.H.L. and C.R.; **Validation,** D.H.L., C.R., and F.P.; **Writing-Original Draft,** D.H.L., C.R., and F.P.; **Writing-Review & Editing,** all other authors commented and agreed on the manuscript; **Visualization,** D.H.L. and C.R.

## Acknowledgments

We thank Dr. Antonio Molina at the technical university of Madrid for providing Col-0^AEQ^ transgenic line seeds and Dr. Cecilia Romanò and Dr. Mads Clausen at the Technical University of Denmark for providing the synthetic RG-I heptasaccharide. The RG-I hydrolase and RG-I lyase were a kind gift from Dr. Kristian Bertel Krogh at Novozymes, Denmark. We thank Julia Watzal and Julian Bünting for technical assistance. Finally, we thank Jana Neuhold, Orla Dunne, and Aranxa Caballero at the Vienna Biocenter Core Facilities (VBCF Protech) for help in molecular cloning and protein production, the VBCF Plant Sciences facilities for the plant growth chambers, and NGS facility.

## Supplemental Information

### Supplemental Figures and legends

**Supplemental Figure. 1.**
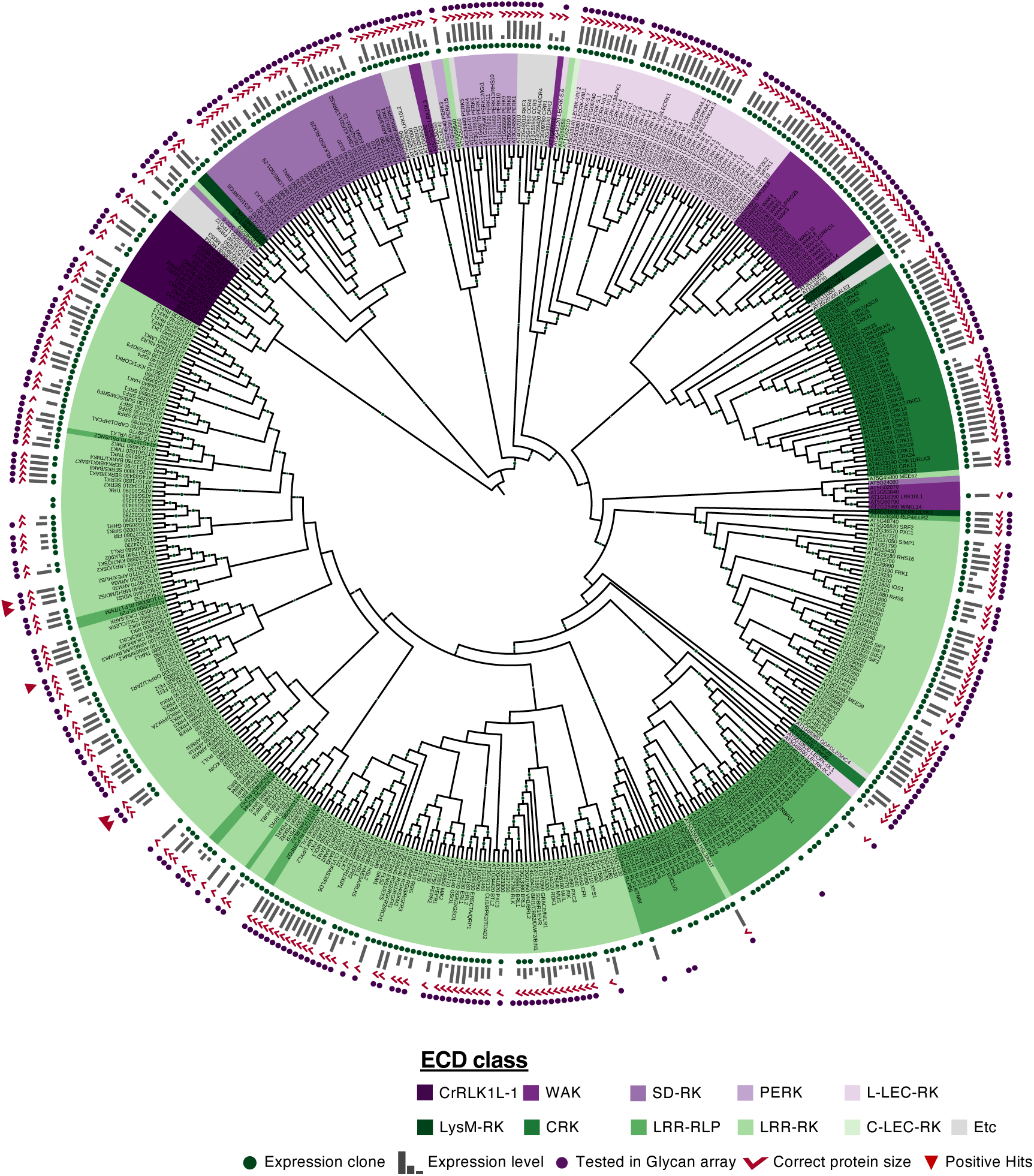
A detailed receptor tree based on ECD sequence similarity of all Arabidopsis plasma membrane-located receptor proteins related to. Fig. 1B. ECD classes were distinguished by different color codes. Bootstrap values above 50 are represented with different dot sizes on each node. Proteins showing larger than expected ECD sizes (Supplemental Table 1) in the immunoblotting assays (Supplemental Fig. 2) are indicated with red check marks.

**Supplemental Figure. 2.**
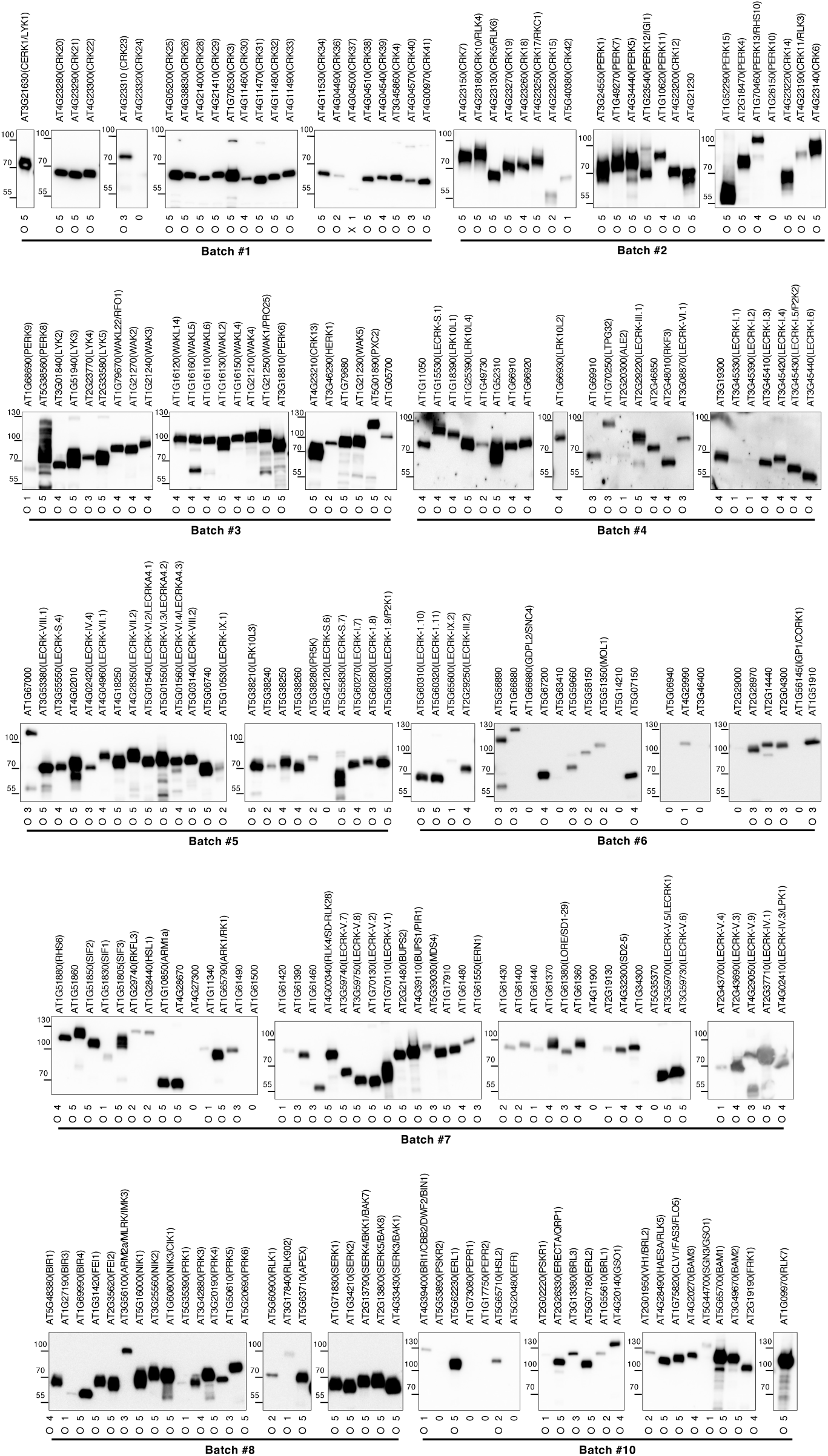
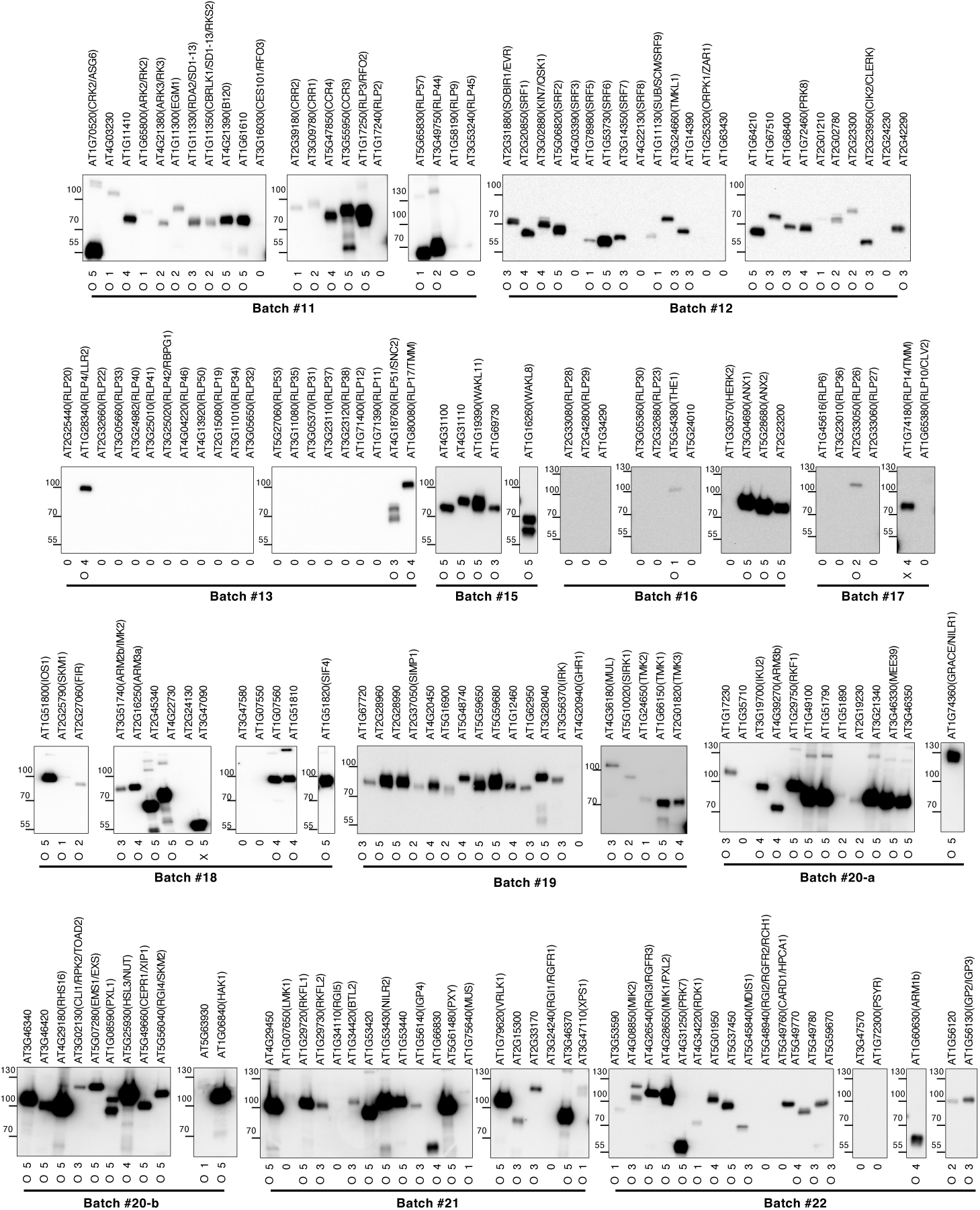
Expression level test of recombinant ECD proteins using immunoblotting. Anti-V5 antibody was used to detect recombinant ECD protein. Gene numbers are indicated at the top of the blots, and expression scores (0-5) are indicated at the bottom. For proteins with detectable expression (scores 1-5), size assessments (O, X) were also provided to evaluate whether the observed molecular weights were consistent with or smaller than the expected sizes, potentially due to protein degradation.

**Supplemental Figure. 3.**
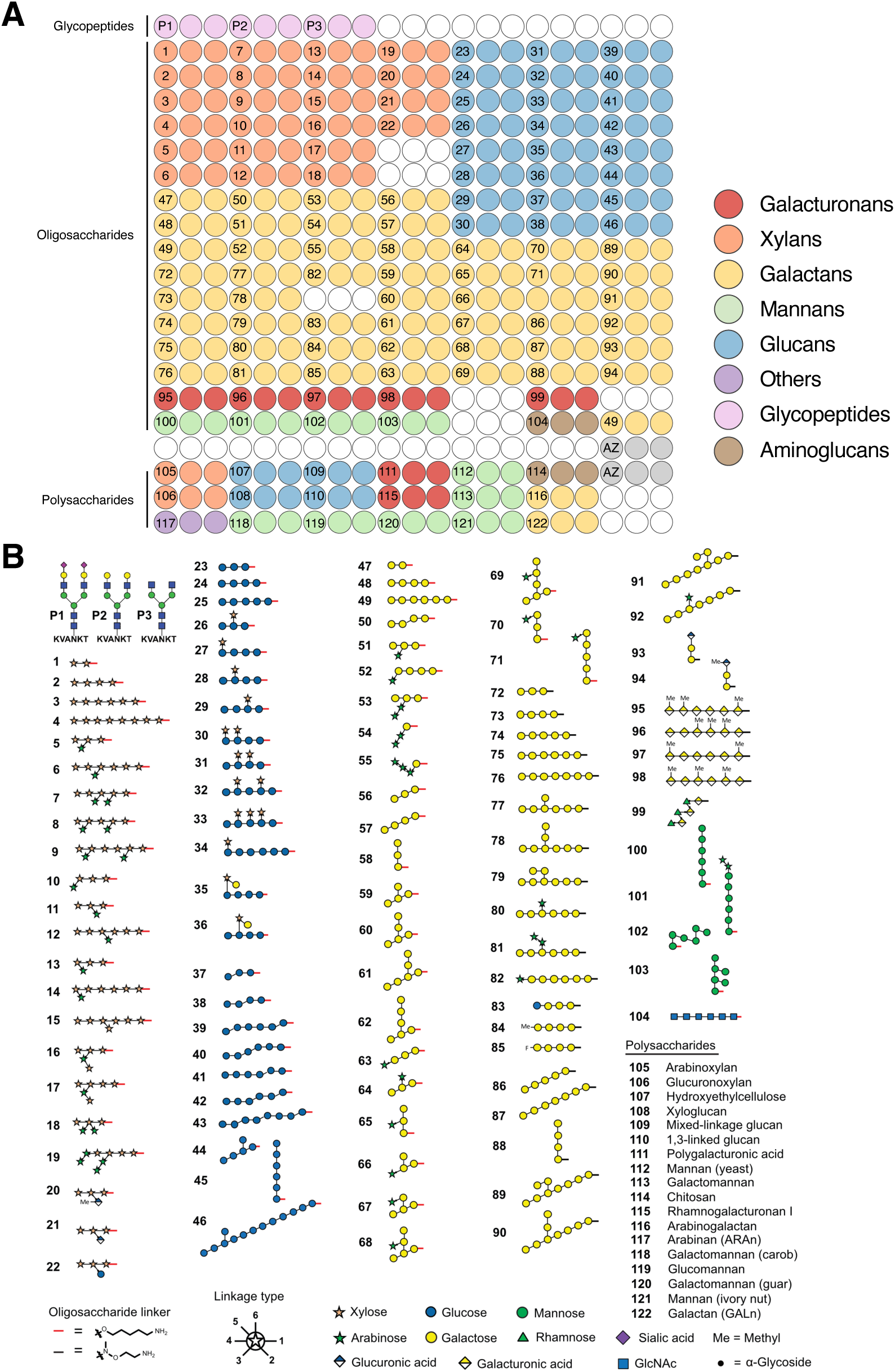
Printing pattern for the plant glycan array. **A.** Printing pattern for each of the 16 fields on a glass slide. **B.** Complete list of printed compounds, including the structures of the oligosaccharides. The angle of the connection between glycan symbols indicates the linkage type, as displayed in the legend below. Methyl indicates methylesterification. The strings for all oligosaccharides are given in Supplemental Table 2. The oligosaccharides were chemically synthesized, the glycopeptides P1-P3 were purified from egg yolk. The sources of the purchased polysaccharides are indicated in Supplemental Table 2.

**Supplemental Figure. 4.**
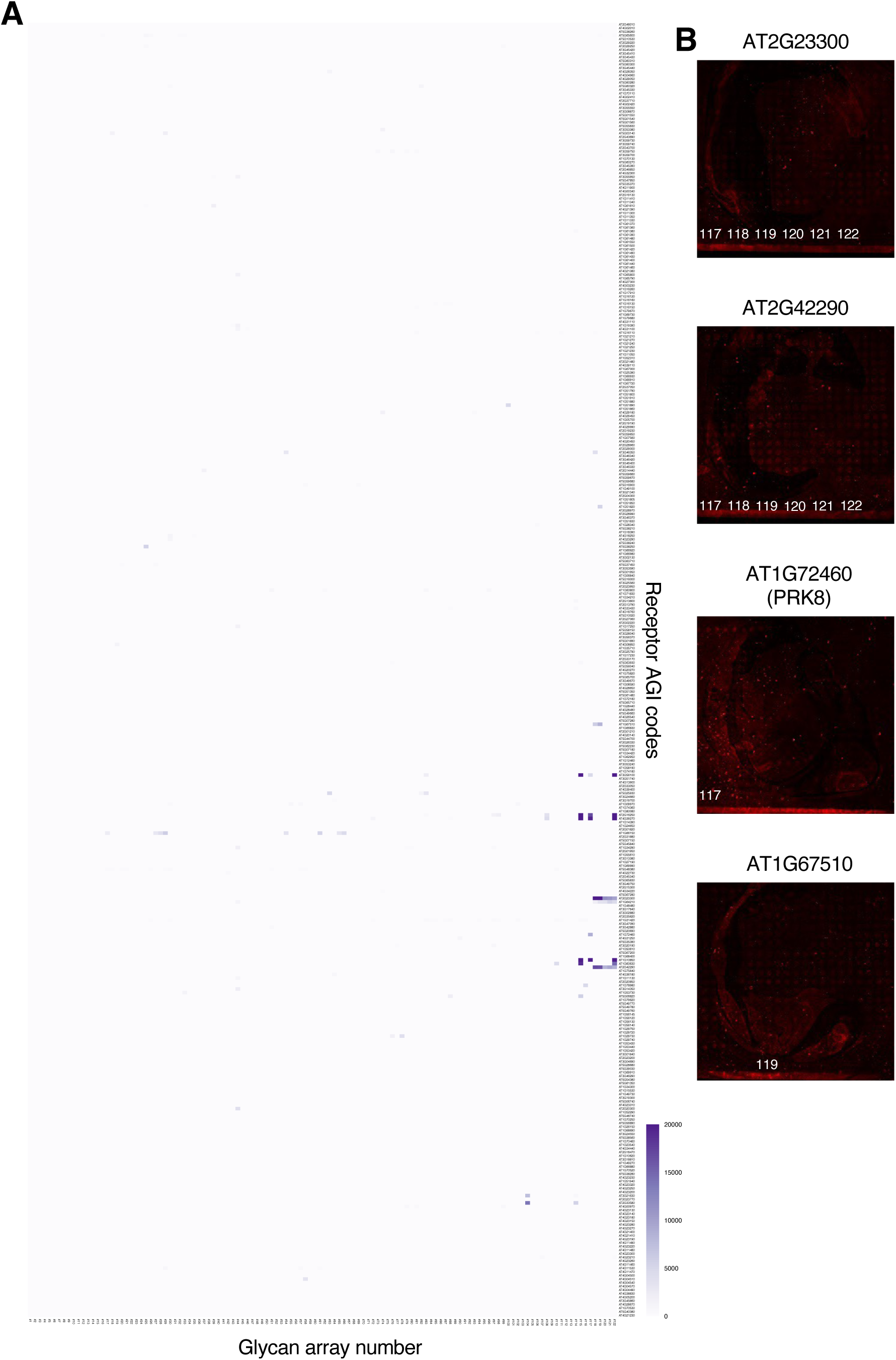
Heatmap of glycan array fluorescence intensity and representative array images showing false positive signals. **A.** Heatmap of all glycan array fluorescence intensities across 357 tested receptor ECDs and 122 glycans. Background-corrected and normalized fluorescence values from Supplemental Table 2 were used for the heatmap. Data points with a coefficient of variation above 60% or median absolute deviation below 0.75× were excluded as they were considered unreliable data points. The color scale shows normalized mean fluorescence intensities from 0 to 20,000. Receptor AGI codes are listed on the right, and glycan numbers printed on the array are shown at the bottom. **B.** Representative scans of 4 ECDs (AT2G23300, AT2G42290, AT1G72460, and AT1G67510) that showed fluorescence signals above the CERK1 values (7,430) in addition to ARM^ECD^s. These fluorescence signals were determined to be false positives because these ECDs were tested on the same slide, and on this slide, the lower border of the applied grid was positioned very close to the last printed row. This led to relatively high fluorescence values in some cases due to interference from the grid border rim, and these signals were therefore considered false positives upon visual inspection.

**Supplemental Figure. 5.**
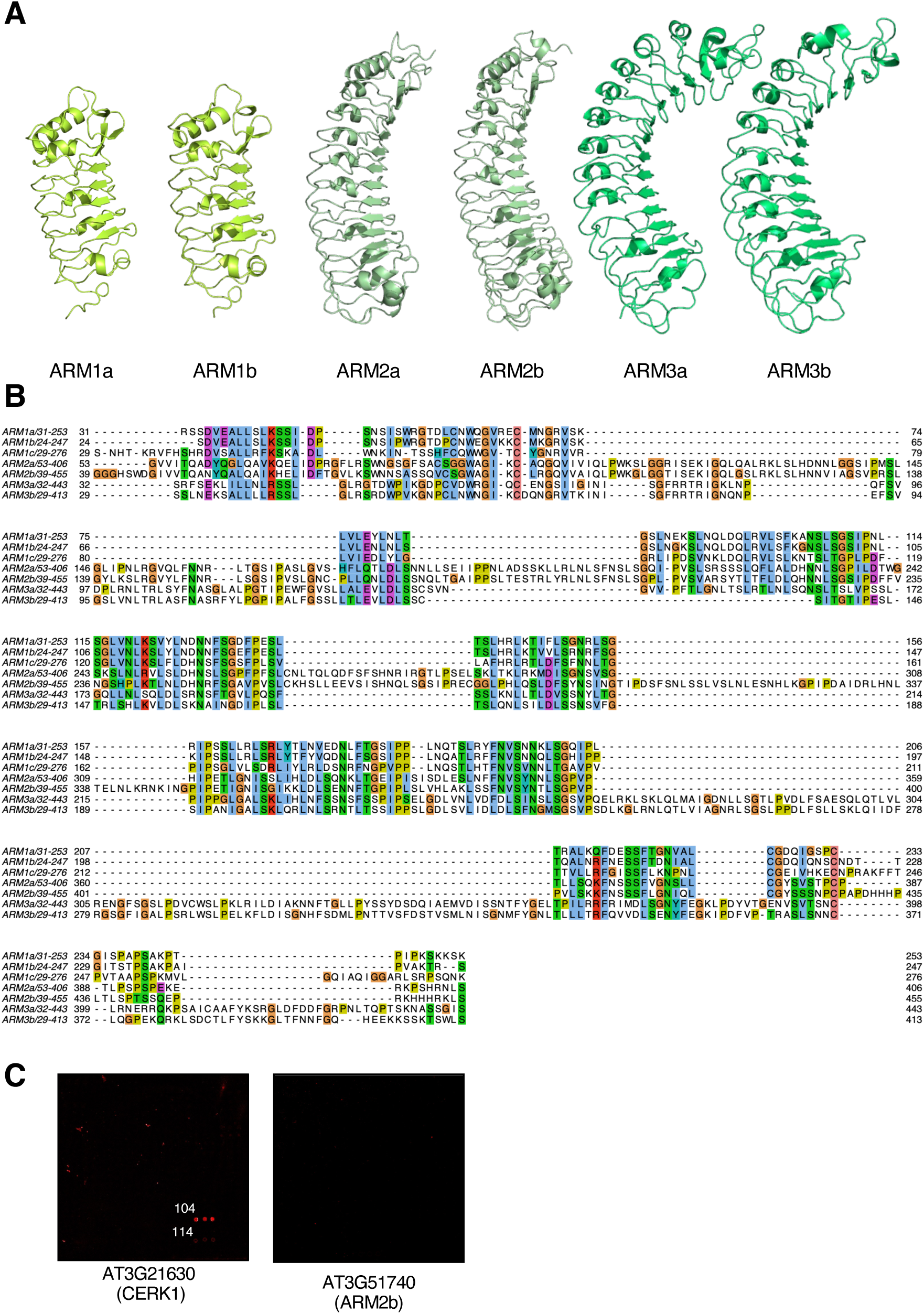
Predicted ARM^ECD^ structures and sequence alignments. **A.** ECD regions from AlphaFold2-predicted structures of ARM^ECD^ proteins with mean pLDDT confidence scores: ARM1a (87.7), ARM1b (87.5), ARM2a (90.0), ARM2b (88.3), ARM3a (91.8), and ARM3b (89.7). **B.** Multiple sequence alignment of ARM^ECD^s. Alignment generated using MAFFT-L-INS-i (v7.511) and displayed with Jalview (ver 2.11.4). Residues are colored according to the Clustal color scheme, based on amino acid properties. Numbers indicate sequence positions. **D.** Interaction assay results from the chitin elicitor receptor kinase 1 (CERK1) and ARM2b in the glycan array screening.

**Supplemental Figure. 6.**
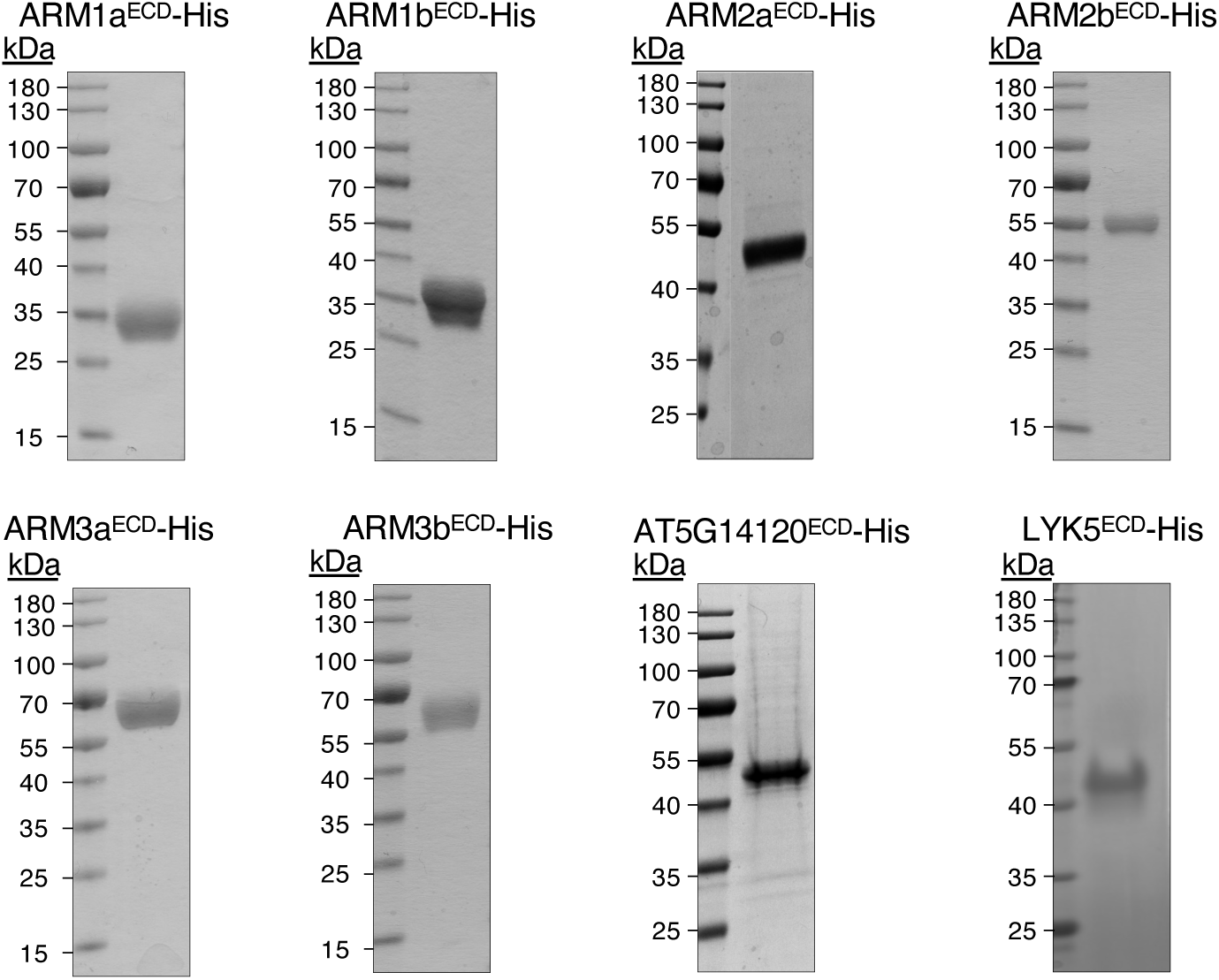
SDS-PAGE gel stained with EZBlue Gel staining reagent showing purified recombinant ECD proteins for MST assay.

**Supplemental Figure. 7.**
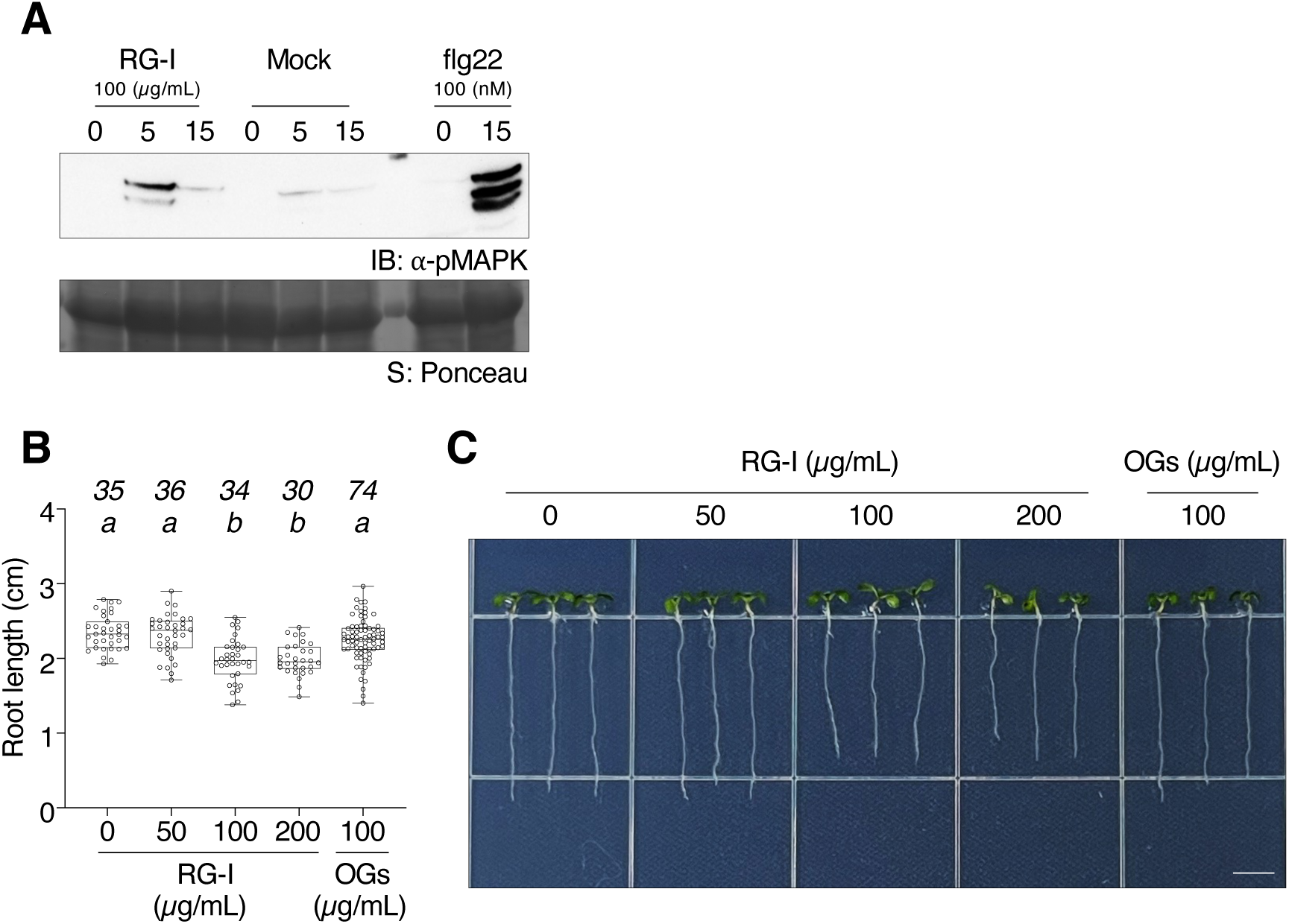
RG-I treatment induces slight root growth inhibition and MAPK activation. **A.** Immunoblotting (IB) analysis of MAPK phosphorylation in wild-type (WT) Arabidopsis seedlings treated with Mock, RG-I or flg22. Treatment conditions, including concentrations and durations, are indicated at the top of the blots. This experiment was repeated with a similar pattern. Equal loading of protein samples on the blot was controlled Coomassie Brilliant blue (C.B.B.) (bottom), which visualizes the Rubisco large subunit (rbcL). **B.** Root growth inhibition rate was assessed by measuring root length 6 days after germination upon treatment with either Mock, OGs, or various concentrations of RG-I. Letters above the boxes (a-b) indicate the results of a one-way ANOVA followed by a Tukey’s multiple comparison test. Groups with the same letter are indistinguishable at >95% confidence. Numbers of biologically independent observations (n) are indicated above the letters. **C.** A representative picture of seedlings for each treatment is shown at the top. Scale bar, 5 mm.

**Supplemental Figure. 8.**
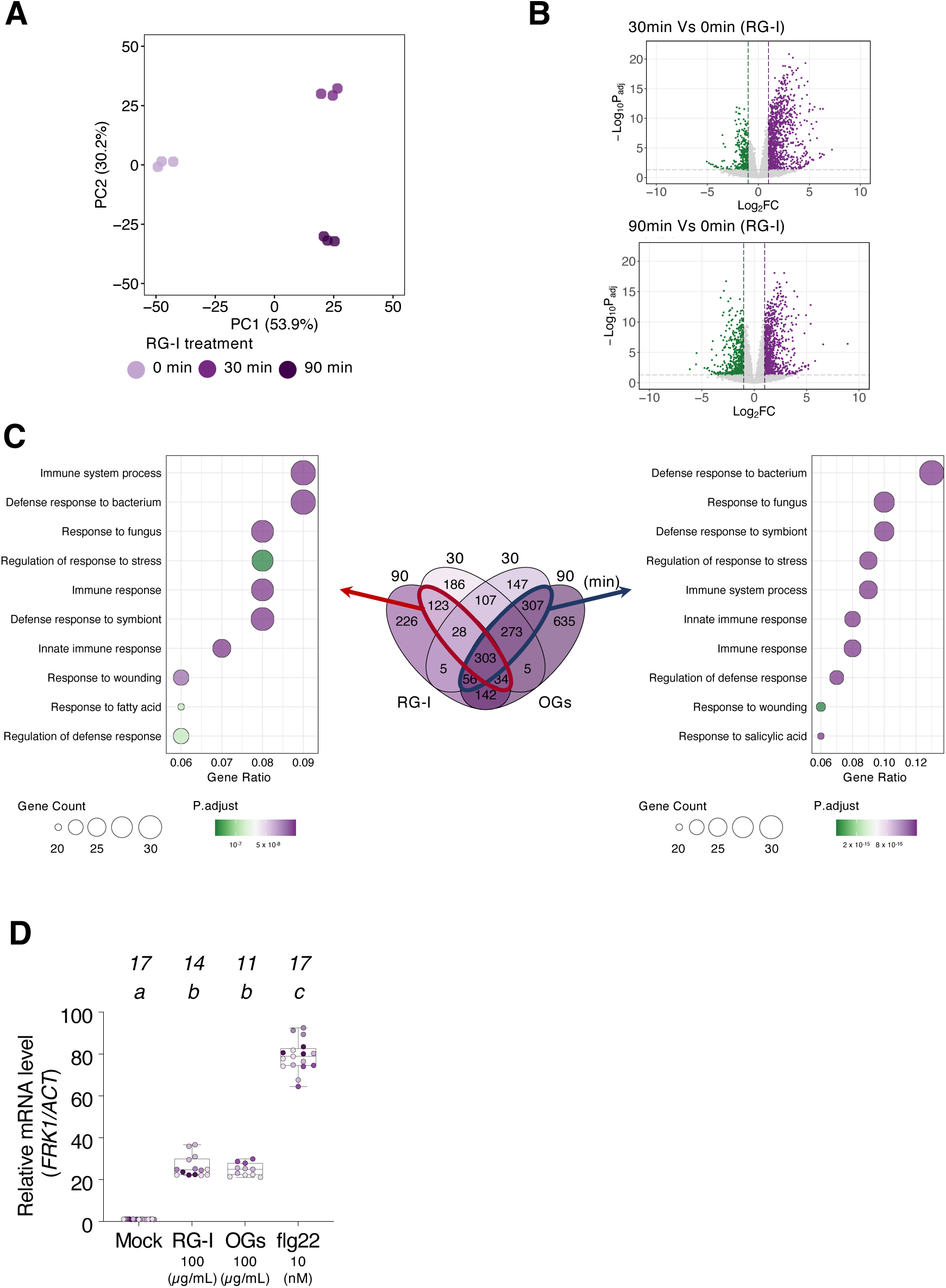
Detailed RNAseq analysis of Arabidopsis seedlings upon RG-I or OGs (OG^DP10-15^) treatment. **A.** Principal component analysis (PCA) performed on the nine samples, including Arabidopsis seedlings treated with RG-I at 0 min, 30 min, and 90 min. **B.** Volcano plots showing RNAseq results comparing 30 min vs 0 min (top) and 90 min vs 0 min (bottom) following RG-I treatment. **C.** Gene Ontology (GO) analysis showing enrichment in biological process categories from overlapping upregulated DEGs by either at both 30 min and 90 min following RG-I treatment or OGs treatment. Bubble chart indicates the calculated gene ratio in each GO term with adjusted P-values. **D.** qPCR analysis showing relative expression levels of *FRK1* normalized to *ACTIN2/8*. Values are presented as relative expression levels compared to Mock sample after 2 h treatment with RG-I (100 μg/mL), OGs (100 μg/mL) or flg22 (10 nM). The box plot represents the first and third quartiles, centered by the median, and whiskers extend to the minimum and maximum data points. Numbers of biological replications (*n*) are indicated above the letters, and dots with different colors indicate different experimental trials. Letters above the boxes (a-c) indicate the results of statistical analysis using a linear mixed effect model with one-way ANOVA followed by a Tukey’s multiple comparison test (two-sided, adjusted for multiple comparisons, *p* < 0.05).

**Supplemental Figure. 9.**
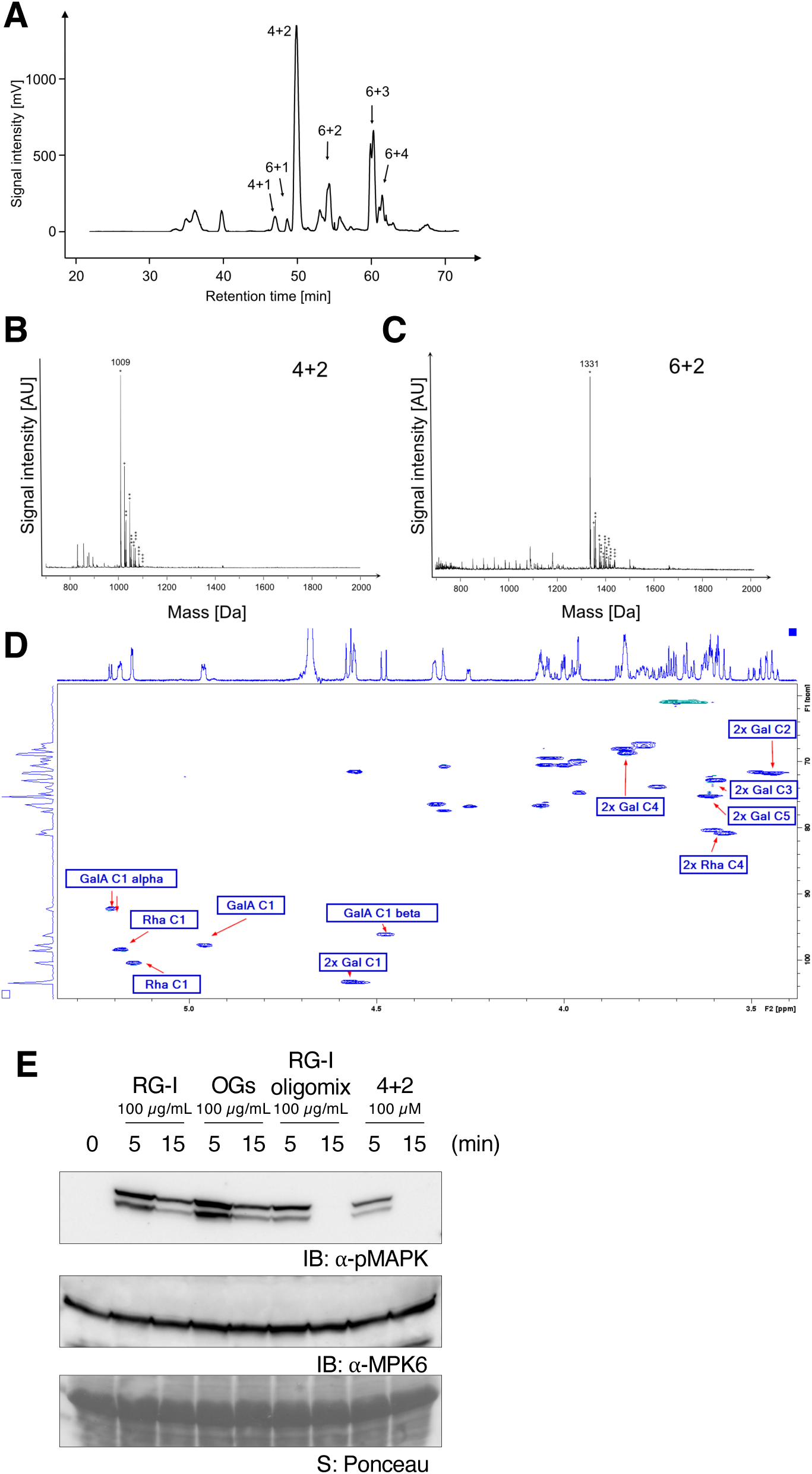
Preparation of RG-I derivate oligosaccharides. **A.** Preparative separation of the RG-I oligosaccharide mixture using a PGC column. Displayed is the chromatogram of the Evaporative Light Scattering Detector (ELSD). **B and C**. MALDI-TOF-MS analyses of the oligosaccharides determined to be 4+2 and 6+2, respectively. The mass of the Na^+^-adduct is indicated. Additional adducts of the same oligosaccharide are labeled with * (Na^+^) and + (K^+^), which correspond to the number of the respective ions in the molecule. Note that H^+^ in GalA in the oligosaccharides may also be replaced with Na^+^ or K^+^. **D.** HSQC Overview of 4+2, relevant anomeric, Gal and Rhamnose signals are flagged. While the mass spectrometric analyses indicated the composition of monosaccharides in the purified RG-I oligosaccharides, the linkages remained unclear. For example, in the oligosaccharide 4+2 (four backbone residues and two attached galactoses), the two galactose units could (i) be attached to two different rhamnose residues or (ii) represent a β1,4-linked digalactoside attached to one of the rhamnose residues. To differentiate between these two scenarios, we performed a full structure elucidation of 4+2. Within this process it became apparent that the anomeric configuration of both Gal residues was equatorial (*J*_H-1,C-1_ = 162 HZ, *J*_H-1,H-2_∼7.9 Hz) and all remaining signals of the two Gal residues were almost indifferentiable from each other in the hydrogen and carbon dimension (C-1=104.02/104.15, C2= 72.43, C3=73.51, C4=69.47, C5=75.91, C6=61.76 ppm). All galactose signals were in good agreement with signals of 1-*O*-methy galactopyranoside^73,74^. These chemical shifts point towards scenario i, since scenario ii would require position 4 or 3 of one Gal residue to be glycosylated, which would significantly shift the signal in both the proton and more importantly the carbon dimension towards higher ppm values^74^. Furthermore, we investigated the connectivity between the Gal residues and their site of attachment on the RG-I backbone-like structure in 4+2. Based on the long range Hetero Multiple Bond Correlation (HMBC) from positions 1 of the Gal residues, we could identify two signals with a characteristic ^13^C-glycosylation shift (81.15 and 81.59 ppm) which were found to belong to the set of rhamnose signals. These two signals were then later identified based on their characteristic triplet coupling pattern (*J*=9.9 Hz) and via Total Correlation Spectroscopy (TOCSY) to be positions 4 of both rhamnose units. We therefore concluded that the proposal for structure I is more plausible. With the information obtained from the 4+2 structure elucidation we investigated the substitution pattern of 6+2. Despite the small sample quantity, we were able to show via overlay of the HSQC and TOCSY spectra that the substitution pattern is similar to the 4+2 structure, since the galactose HSQC cross peaks of 4+2 and 6+2 completely overlay, as do the TOCSY crosspeaks. We therefore conclude that in both structures the galactose residues are located at the backbone of RG-I. **E.** Immunoblotting (IB) analyses of MAPK phosphorylation in wild-type (WT) Arabidopsis seedlings treated with RG-I, OGs, or RG-I fragments. Treatment conditions, including concentrations and durations, are indicated at the top of the blots. This experiment was repeated with a similar pattern. Equal loading of protein samples on the blot was controlled by anti-MPK blotting (middle) and Ponceau staining (bottom), which visualizes the Rubisco large subunit (rbcL).

**Supplemental Figure. 10.**
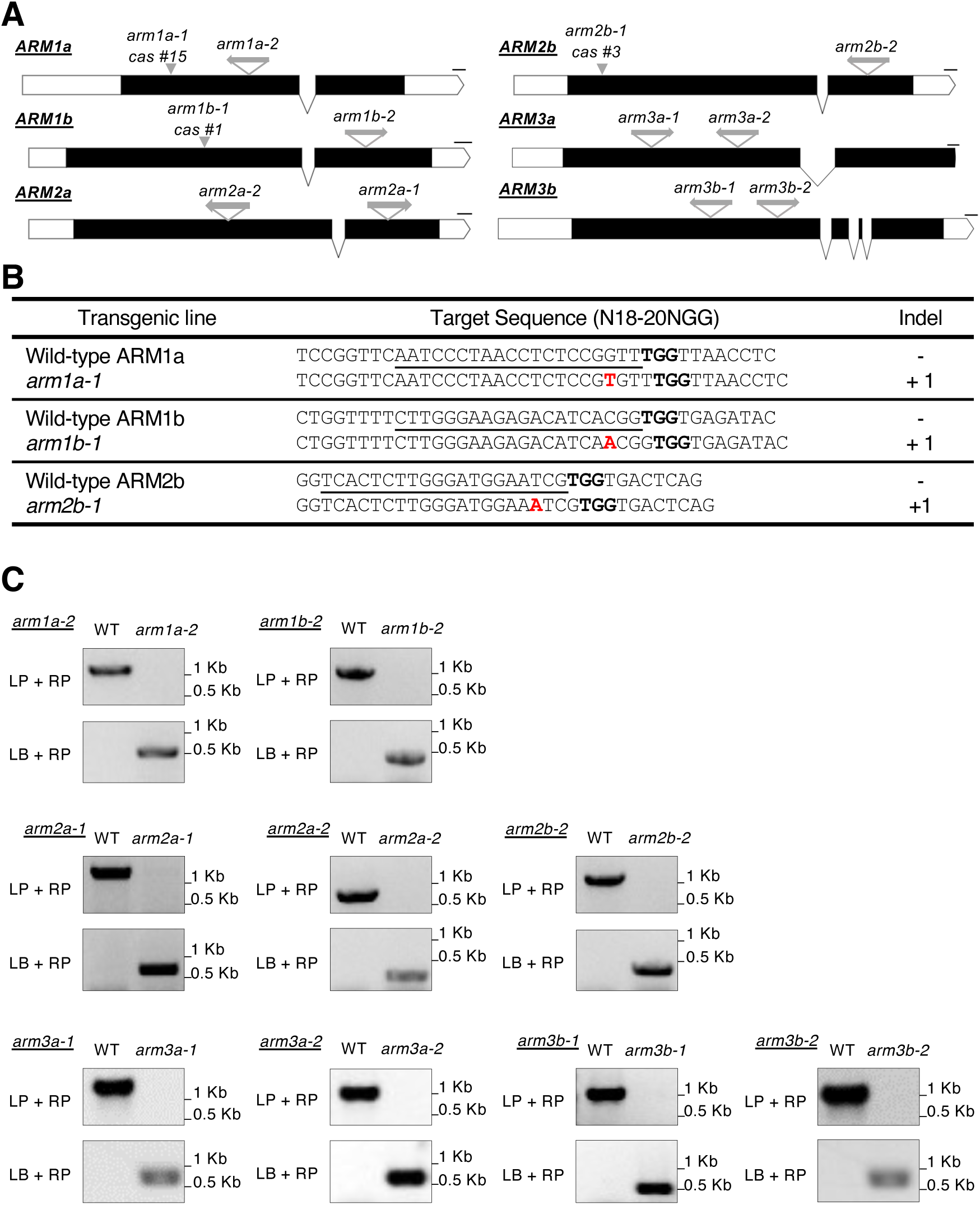
Genotyping and mutation analysis of *ARM* single mutants. **A.** Schematic representation showing T-DNA insertion or gene editing locations within *ARM* gene structure. **B.** Gene-editing mutations identified in *ARM1a*, *ARM1b*, and *ARM2b* corresponding to *arm1a-1*, *arm1b-1*, and *arm2b-1* mutants, respectively. **C.** PCR-based genotyping results of ARM single T-DNA mutants.

**Supplemental Figure. 11.**
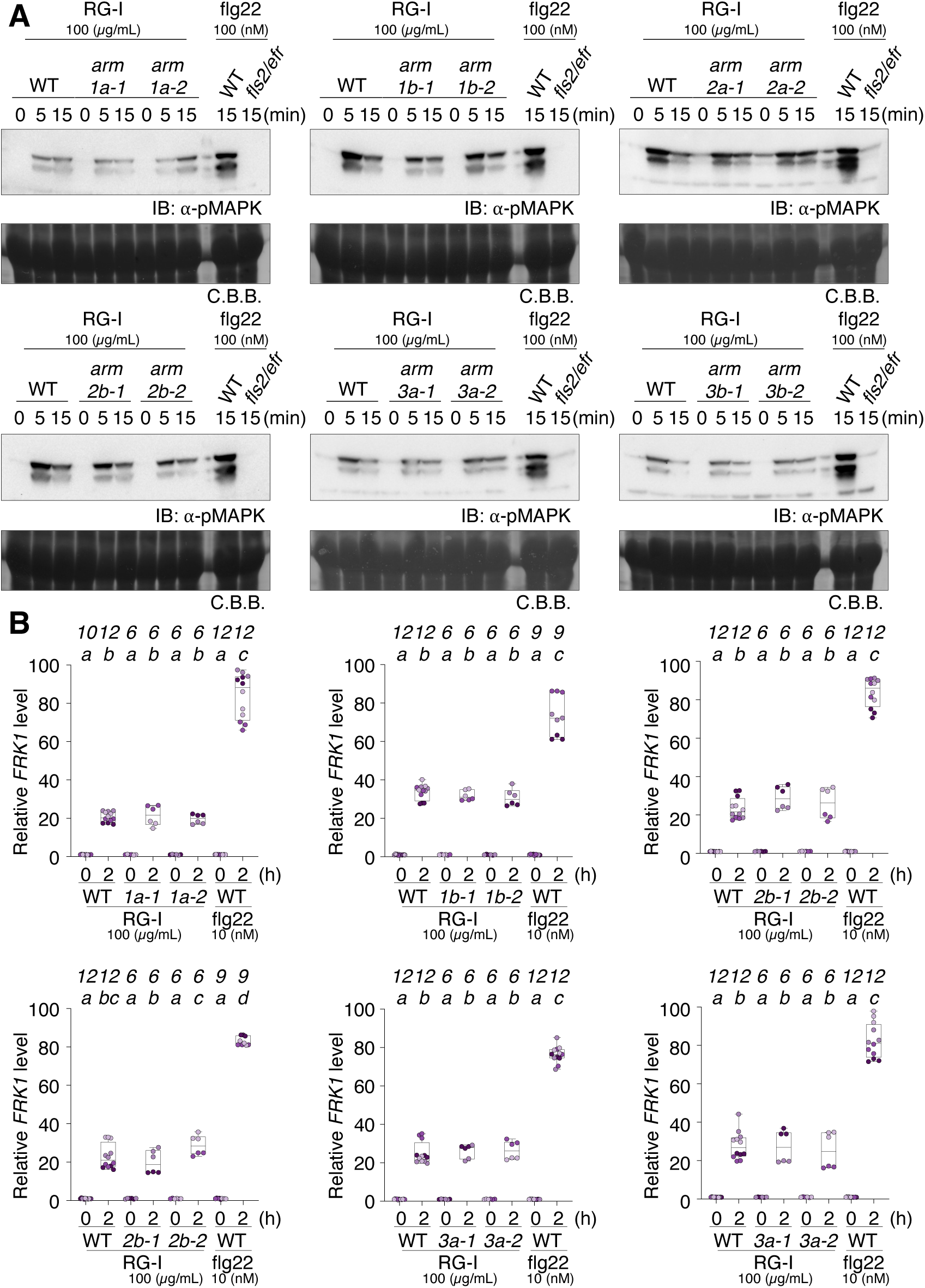
PTI responses in ARM single mutants. **A.** Immunoblotting (IB) analyses of MAPK phosphorylation in wild-type (WT) Arabidopsis seedlings and ARM single mutant Arabidopsis seedlings with RG-I treatment. Treatment conditions, including concentrations and durations, are indicated at the above the blots. This experiment was repeated twice with similar results. Equal loading of protein samples on the blot was controlled by Coomassie Brilliant blue (C.B.B.) (bottom), which visualizes the Rubisco large subunit (rbcL). **B.** qPCR analysis showing relative expression levels of *FRK1* normalized to *ACTIN2/8*. Values are presented as relative expression levels compared to each genotype’s corresponding untreated (0 h) sample after 2 h treatment with RG-I (100 μg/mL) or flg22 (10 nM). The box plot represents the first and third quartiles, centered by the median, and whiskers extend to the minimum and maximum data points. Numbers of biological replications (*n*) are indicated above the letters, and dots with different colors indicate different experimental trials. Letters above the boxes (a-d) indicate the results of statistical analysis using a linear mixed effect model with one-way ANOVA followed by a Tukey’s multiple comparison test (two-sided, adjusted for multiple comparisons, *p* < 0.05).

**Supplemental Figure. 12.**
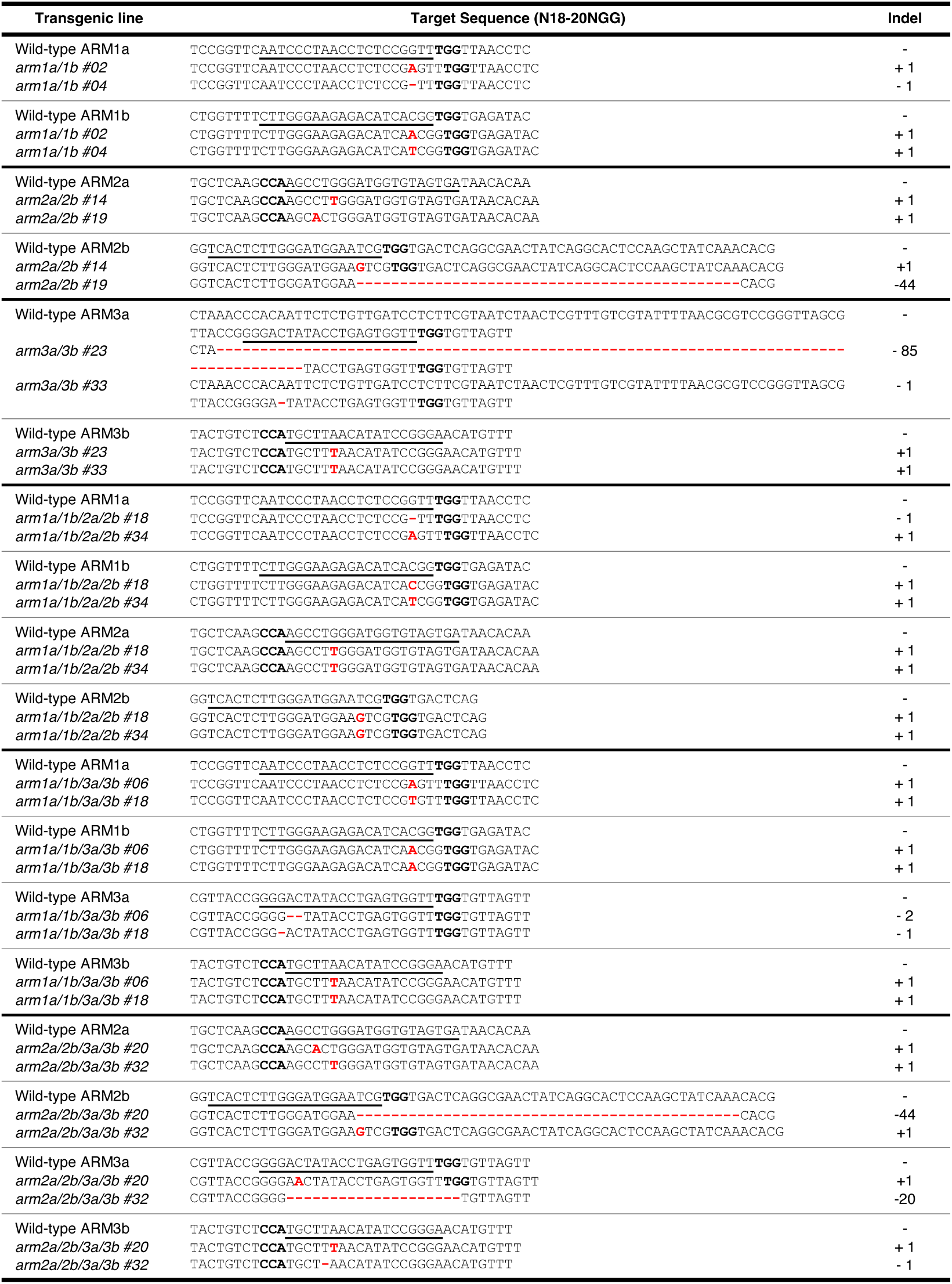
Gene-editing mutations identified in *ARM* genes from double mutants and quadruple mutants.

**Supplemental Figure. 13.**
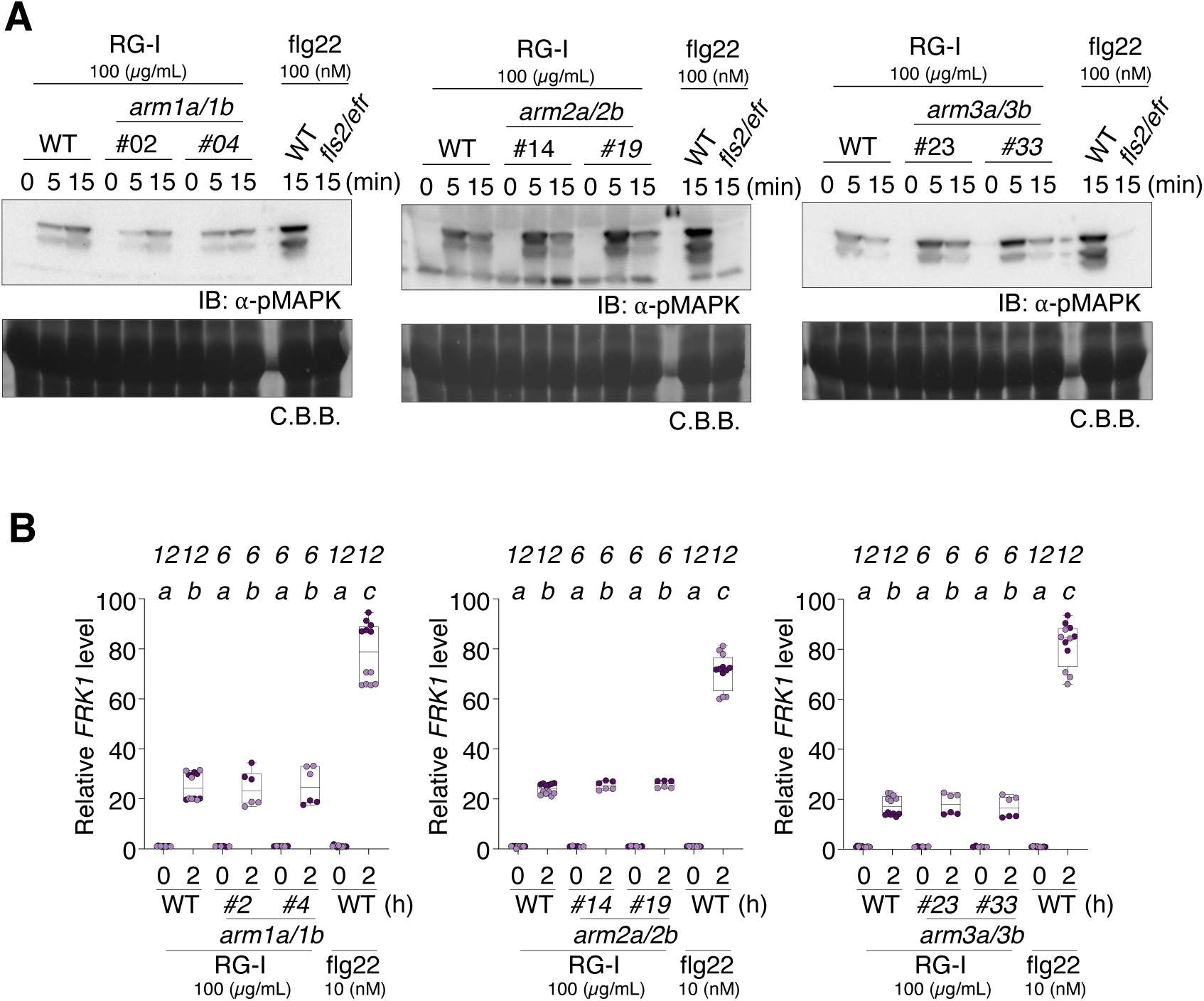
PTI responses in ARM double mutants. **A.** Immunoblotting (IB) analyses of MAPK phosphorylation in wild-type (WT) Arabidopsis seedlings and ARM double mutant Arabidopsis seedlings with RG-I treatment. Treatment conditions, including concentrations and durations, are indicated at the above the blots. This experiment was repeated twice with similar results. Equal loading of protein samples on the blot was controlled by Coomassie Brilliant blue (C.B.B.) (bottom), which visualizes the Rubisco large subunit (rbcL). **B.** qPCR analysis showing relative expression levels of *FRK1* normalized to *ACTIN2/8*. Values are presented as relative expression levels compared to each genotype’s corresponding untreated (0 h) sample after 2 h treatment with RG-I (100 μg/mL) or flg22 (10 nM). The box plot represents the first and third quartiles, centered by the median, and whiskers extend to the minimum and maximum data points. Numbers of biological replications (*n*) are indicated above the letters, and dots with different colors indicate different experimental trials. Letters above the boxes (a-d) indicate the results of statistical analysis using a linear mixed effect model with one-way ANOVA followed by a Tukey’s multiple comparison test (two-sided, adjusted for multiple comparisons, *p* < 0.05).

**Supplemental Figure. 14.**
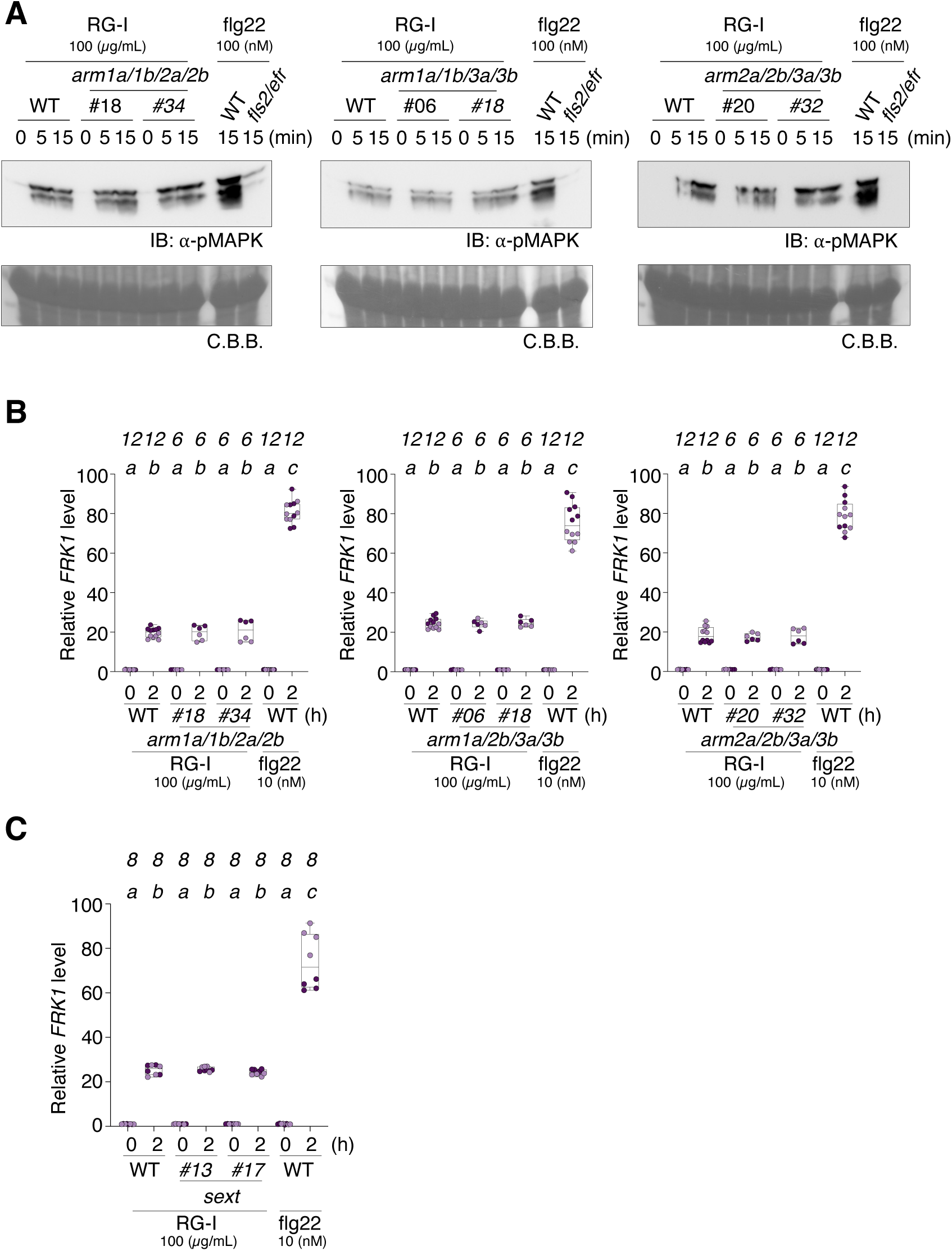
PTI responses in ARM quadruple and sextuple mutants. **A.** Immunoblotting (IB) analyses of MAPK phosphorylation in wild-type (WT) Arabidopsis seedlings and ARM quadruple mutant Arabidopsis seedlings with RG-I treatment. Treatment conditions, including concentrations and durations, are indicated at the above the blots. This experiment was repeated twice with similar results. Equal loading of protein samples on the blot was controlled by Coomassie Brilliant blue (C.B.B.) (bottom), which visualizes the Rubisco large subunit (rbcL). **B and C.** qPCR analysis showing relative expression levels of *FRK1* normalized to *ACTIN2/8*. Values are presented as relative expression levels compared to each genotype’s corresponding untreated (0 h) sample after 2 h treatment with RG-I (100 μg/mL) or flg22 (10 nM). The box plot represents the first and third quartiles, centered by the median, and whiskers extend to the minimum and maximum data points. Numbers of biological replications (*n*) are indicated above the letters, and dots with different colors indicate different experimental trials. Letters above the boxes (a-d) indicate the results of statistical analysis using a linear mixed effect model with one-way ANOVA followed by a Tukey’s multiple comparison test (two-sided, adjusted for multiple comparisons, *p* < 0.05).

**Supplemental Figure. 15.**
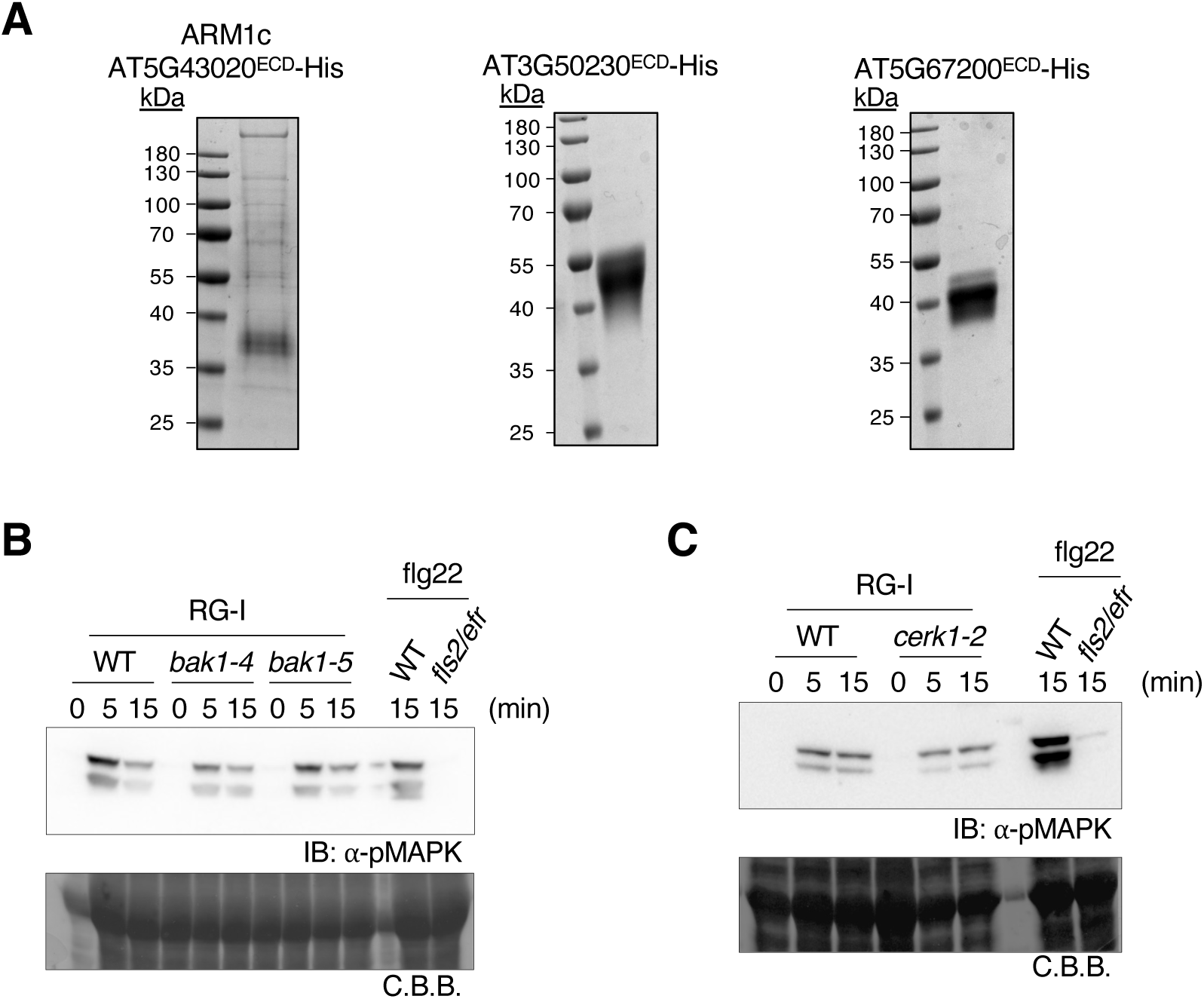
Purified ECD proteins from ARM1-containing clades from the receptor tree and MAPK activations in well-studied co-receptor mutants. **A.** SDS-PAGE gel stained with EZBlue Gel staining reagent showing purified recombinant ECD proteins for MST assay. **B and C.** Immunoblotting (IB) analyses of MAPK phosphorylation in wild-type (WT) Arabidopsis seedlings and either *bak1* mutants (B) or *cerk1* mutant (C) Arabidopsis seedlings with RG-I treatment. Treatment conditions, including concentrations and durations, are indicated at the above the blots. This experiment was repeated twice with similar results. Equal loading of protein samples on the blot was controlled by Coomassie Brilliant blue (C.B.B.) (bottom), which visualizes the Rubisco large subunit (rbcL).

**Supplemental Figure. 16.**
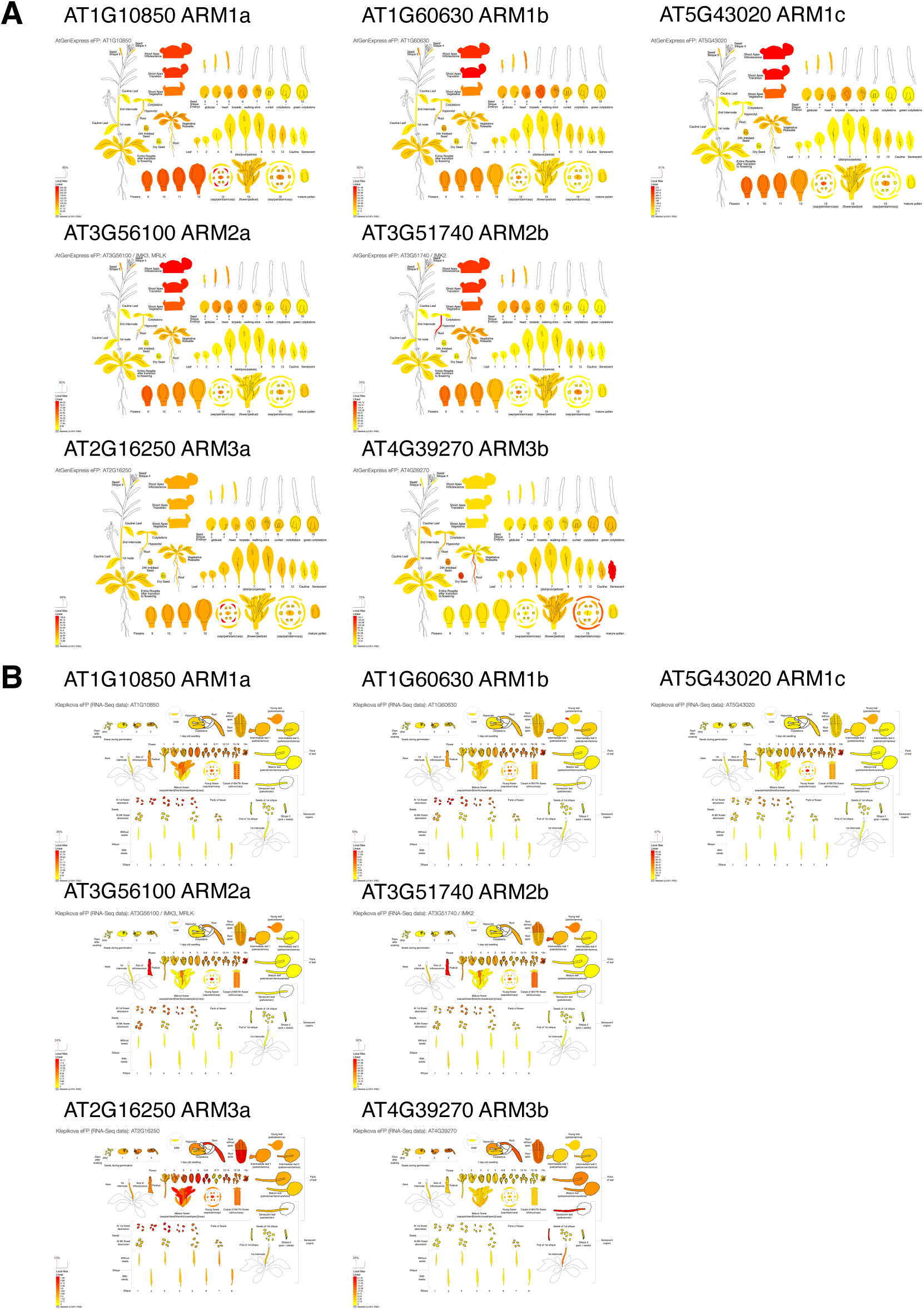
Tissue-specific gene expression patterns of ARM receptors. Expression patterns of each ARM receptor were generated using the AtGeneExpress eFP Browser (https://bar.utoronto.ca/eplant/,^75^) based on publicly available gene expression data.

### Supplemental Tables and legends

**Supplemental Table 1. Summary of Arabidopsis RKs and RLPs analyzed in this study.** Tree_order: Order of receptors in the receptor tree shown in Supplemental Fig. 1. Gene: AGI code. Protein: Representative cDNA identifier. Protein_name: Currently recognized receptor name. Receptor_structure: Type (RLK/RLP) and Class including Leucine-rich repeat (LRR), Wall-associated kinase (WAK), WAK like (WAKL), L-type lectin (L-LEC), C-type lectin (C-LEC), S-domain (SD), Catharanthus roseus RLK (CrRLK), Proline-rich extensin-like RLK (PERK), Cystein-rich RLK (CrRLK), Domain of unknown function 26 (DUF26). DeepTMHMM_results: Amino acid positions for signal peptide, ECD start, ECD end, transmembrane domain (TM), and cytoplasmic domain predicted by DeepTMHMM. Protein_size: Predicted molecular weight (MW) in kDa. MW_(ECD+FC): Predicted molecular weight of extracellular domain (ECD) fused with fragment crystallizable region from antibody heavy chains (FC) domain. Cloning_information: Internal code (Vienna BioCentor) or AddGene repository number. Expression: Batch number for expression experiments; Expression_level represents immunoblotting scores (0-5) from Supplemental Fig. 2; Correct_protein_size marked as “O” indicates observed molecular weight was within expected range or slightly larger than predicted MW_FC. Glycan_array_test: “O” indicates inclusion in glycan array screening. Positive_hit: “O” indicates receptors having more than one value higher than CERK1-chitohexaose threshold.

**Supplemental Table 2. Glycan array experimental data and compound information. All_compund_information** (Sheet 1): Information for all compounds used in the glycan array. For chemically synthesized oligosaccharides: Web universal representation of carbohydrate structures (WURCS) and Glycan workbench structure (GWS) strings. For commercially available polysaccharides: compound names and supplier information. **Normalized_raw_data** (Sheet 2): Background-corrected fluorescence values normalized against negative control spots (printed azide) and adjusted using standard deviation for all 357 receptor ECDs testing against 122 glycans. Values represent [Normalized_Mean] – [Normalized_SD] from measurements (n=3) obtained using PMT800 setting. ARM_detailed_data (Sheet 3): Detailed raw replicate data (Rep1, Rep2, Rep3) with mean and standard deviation values for ARM receptor ECDs. Data obtained using PMT500 setting to optimize signal detection and reduce background noise levels. Mean represents the average of replicates (n=3); Stdev represents standard deviation.

**Supplemental Table 3. DEGs used to analyze GO term (Padj < 0.1 and |log2FC| > 1)** Differentially expressed genes (DEGs) used for GO analysis shown in Fig. 3G (RG-I_OGs_both_Up) and Supplemental Fig. 8C (RG-I_30m_90m_Up and OGs_30m_90m_Up). Statistical significance (adjusted p-value) was determined using the DESeq2 package with a two-sided test for multiple comparisons, applying cut-offs at an adjusted p-value < 0.1 and |log2(Fold Change)| > 1.

**Supplemental Table 4. Primers used in this study.**

## References

Aziz, A., Gauthier, A., Bézier, A., Poinssot, B., Joubert, J.-M., Pugin, A., Heyraud, A., and Baillieul, F. (2007). Elicitor and resistance-inducing activities of β-1,4 cellodextrins in grapevine, comparison with β-1,3 glucans and α-1,4 oligogalacturonides. Journal of Experimental Botany 5810.1093/jxb/erm008.

Bacete, L., Mélida, H., Miedes, E., and Molina, A. (2018). Plant cell wall-mediated immunity: cell wall changes trigger disease resistance responses. The Plant Journal 93:614–636. 10.1111/tpj.13807.

Bacete, L., Mélida, H., Pattathil, S., Hahn, M.G., Molina, A., and Miedes, E. (2017). Plant Pattern Recognition Receptors, Methods and Protocols. Methods in Molecular Biology 1578:13–23. 10.1007/978-1-4939-6859-6_2.

Bartetzko, M.P., Schuhmacher, F., Seeberger, P.H., and Pfrengle, F. (2017). Determining Substrate Specificities of β1,4-Endogalactanases Using Plant Arabinogalactan Oligosaccharides Synthesized by Automated Glycan Assembly. J Org Chem 82:1842–1850. 10.1021/acs.joc.6b02745.

Bartetzko, M.P., Schuhmacher, F., Hahm, H.S., Seeberger, P.H., and Pfrengle, F. (2015). Automated Glycan Assembly of Oligosaccharides Related to Arabinogalactan Proteins. Org Lett 17:4344–4347. 10.1021/acs.orglett.5b02185.

Bigeard, J., Colcombet, J., and Hirt, H. (2015). Signaling mechanisms in pattern-triggered immunity (PTI). Mol Plant 8:521–539. 10.1016/j.molp.2014.12.022.

Bjornson, M., Pimprikar, P., Nürnberger, T., and Zipfel, C. (2021). The transcriptional landscape of Arabidopsis thaliana pattern-triggered immunity. Nature Plants 7:579–586. 10.1038/s41477-021-00874-5.

Brutus, A., Sicilia, F., Macone, A., Cervone, F., and Lorenzo, G.D. (2010). A domain swap approach reveals a role of the plant wall-associated kinase 1 (WAK1) as a receptor of oligogalacturonides. Proceedings of the National Academy of Sciences 107:9452–9457. 10.1073/pnas.1000675107.

Cao, Y., Liang, Y., Tanaka, K., Nguyen, C.T., Jedrzejczak, R.P., Joachimiak, A., and Stacey, G. (2014). The kinase LYK5 is a major chitin receptor in Arabidopsis and forms a chitin-induced complex with related kinase CERK1. eLife 3:e03766. 10.7554/elife.03766.

Choi, J., Tanaka, K., Cao, Y., Qi, Y., Qiu, J., Liang, Y., Lee, S.Y., and Stacey, G. (2014). Identification of a Plant Receptor for Extracellular ATP. Science 343:290–294. 10.1126/science.343.6168.290.

Clasen, S.J., Bell, M.E.W., Borbón, A., Lee, D.-H., Henseler, Z.M., Cuesta-Zuluaga, J.d.l., Parys, K., Zou, J., Wang, Y., Altmannova, V., et al. (2023). Silent recognition of flagellins from human gut commensal bacteria by Toll-like receptor 5. Science immunology 8:eabq7001. 10.1126/sciimmunol.abq7001.

Claverie, J., Balacey, S., Lemaître-Guillier, C., Brulé, D., Chiltz, A., Granet, L., Noirot, E., Daire, X., Darblade, B., Héloir, M.-C., et al. (2018). The Cell Wall-Derived Xyloglucan Is a New DAMP Triggering Plant Immunity in Vitis vinifera and Arabidopsis thaliana. Frontiers in Plant Science 9:1725. 10.3389/fpls.2018.01725.

Côté, F., and Hahn, M.G. (1994). Oligosaccharins: structures and signal transduction. Plant Mol Biol 26:1379–1411. 10.1007/BF00016481.

Dallabernardina, P., Schuhmacher, F., Seeberger, P.H., and Pfrengle, F. (2016). Automated glycan assembly of xyloglucan oligosaccharides. Org Biomol Chem 14:309–313. 10.1039/c5ob02226f.

Dallabernardina, P., Schuhmacher, F., Seeberger, P.H., and Pfrengle, F. (2017). Mixed-Linkage Glucan Oligosaccharides Produced by Automated Glycan Assembly Serve as Tools To Determine the Substrate Specificity of Lichenase. Chemistry 23:3191–3196. 10.1002/chem.201605479.

DeFalco, T.A., and Zipfel, C. (2021). Molecular mechanisms of early plant pattern-triggered immune signaling. Molecular Cell 81:3449–3467. 10.1016/j.molcel.2021.07.029.

Delmer, D., Dixon, R.A., Keegstra, K., and Mohnen, D. (2024). The plant cell wall-dynamic, strong, and adaptable-is a natural shapeshifter. Plant Cell 36:1257–1311. 10.1093/plcell/koad325.

Denoux, C., Galletti, R., Mammarella, N., Gopalan, S., Werck, D., De Lorenzo, G., Ferrari, S., Ausubel, F.M., and Dewdney, J. (2008). Activation of defense response pathways by OGs and Flg22 elicitors in Arabidopsis seedlings. Mol Plant 1:423–445. 10.1093/mp/ssn019.

Du, J., Anderson, C.T., and Xiao, C. (2022). Dynamics of pectic homogalacturonan in cellular morphogenesis and adhesion, wall integrity sensing and plant development. Nature Plants 8:332–340. 10.1038/s41477-022-01120-2.

Gómez-Gómez, L., Felix, G., and Boller, T. (1999). A single locus determines sensitivity to bacterial flagellin in Arabidopsis thaliana. Plant J 18:277–284. 10.1046/j.1365-313x.1999.00451.x.

Hallgren, J., Tsirigos, K.D., Pedersen, M.D., Armenteros, J.J.A., Marcatili, P., Nielsen, H., Krogh, A., and Winther, O. (2022). DeepTMHMM predicts alpha and beta transmembrane proteins using deep neural networks. bioRxiv 10.1101/2022.04.08.487609.

Harholt, J., Suttangkakul, A., and Vibe Scheller, H. (2010). Biosynthesis of pectin. Plant Physiol 153:384–395. 10.1104/pp.110.156588.

He, Z., Webster, S., and He, S.Y. (2022). Growth-defense trade-offs in plants. Curr Biol 32:R634–R639. 10.1016/j.cub.2022.04.070.

Herold, L., Hua, C., Kohorn, B., Nürnberger, T., DeFalco, T., and Zipfel, C. (2024). Arabidopsis WALL-ASSOCIATED KINASES are not required for oligogalacturonide-induced signaling and immunity. bioRxiv 10.1101/2024.04.15.589471.

Hématy, K., Cherk, C., and Somerville, S. (2009). Host-pathogen warfare at the plant cell wall. Curr Opin Plant Biol 12:406–413. 10.1016/j.pbi.2009.06.007.

Jing, Y., Zheng, X., Zhang, D., Shen, N., Wang, Y., Yang, L., Fu, A., Shi, J., Zhao, F., Lan, W., et al. (2019). Danger-Associated Peptides Interact with PIN-Dependent Local Auxin Distribution to Inhibit Root Growth in Arabidopsis. Plant Cell 31:1767–1787. 10.1105/tpc.18.00757.

Kaku, H., Nishizawa, Y., Ishii-Minami, N., Akimoto-Tomiyama, C., Dohmae, N., Takio, K., Minami, E., and Shibuya, N. (2006). Plant cells recognize chitin fragments for defense signaling through a plasma membrane receptor. Proceedings of the National Academy of Sciences 103:11086–11091. 10.1073/pnas.0508882103.

Kawai, T., Ikegawa, M., Ori, D., and Akira, S. (2024). Decoding Toll-like receptors: Recent insights and perspectives in innate immunity. Immunity 57:649–673. 10.1016/j.immuni.2024.03.004.

Khosravi, C., Kun, R.S., Visser, J., Aguilar-Pontes, M.V., de Vries, R.P., Battaglia, E., Khosravi, C., Kun, R.S., Visser, J., Aguilar-Pontes, M.V., et al. (2017). In vivo functional analysis of L-rhamnose metabolic pathway in Aspergillus niger: a tool to identify the potential inducer of RhaR. BMC Microbiology 2017 17:1 1710.1186/s12866-017-1118-z.

Klarzynski, O., Plesse, B., Joubert, J.-M., Yvin, J.-C., Kopp, M., Kloareg, B., and Fritig, B. (2000). Linear β-1,3 Glucans Are Elicitors of Defense Responses in Tobacco. Plant Physiology 12410.1104/pp.124.3.1027.

Kohorn, B.D., Greed, B.E., Mouille, G., Verger, S., and Kohorn, S.L. (2021). Effects of Arabidopsis wall associated kinase mutations on ESMERALDA1 and elicitor induced ROS. PLoS One 16:e0251922. 10.1371/journal.pone.0251922.

Kubicek, C.P., Starr, T.L., and Glass, N.L. (2014). Plant cell wall-degrading enzymes and their secretion in plant-pathogenic fungi. Annu Rev Phytopathol 52:427–451. 10.1146/annurev-phyto-102313-045831.

Lee, D.-H., Lee, H.-S., and Belkhadir, Y. (2021). Coding of plant immune signals by surface receptors. Current Opinion in Plant Biology 62:102044. 10.1016/j.pbi.2021.102044.

Lee, D.-H., Lee, H.-S., Choi, M.-S., Parys, K., Honda, K., Kondoh, Y., Lee, J.-M., Edelbacher, N., Heo, G., Enugutti, B., et al. (2024). Reprogramming of flagellin receptor responses with surrogate ligands. Nature Communications 1510.1038/s41467-024-54271-5.

Letunic, I., and Bork, P. (2024). Interactive Tree of Life (iTOL) v6: recent updates to the phylogenetic tree display and annotation tool. Nucleic Acids Res 52:W78–W82. 10.1093/nar/gkae268.

Liu, C., Yu, H., Voxeur, A., Rao, X., and Dixon, R.A. (2023). FERONIA and wall-associated kinases coordinate defense induced by lignin modification in plant cell walls. Sci Adv 9:eadf7714. 10.1126/sciadv.adf7714.

Malinovsky, F.G., Fangel, J.U., and Willats, W.G. (2014). The role of the cell wall in plant immunity. Front Plant Sci 5:178. 10.3389/fpls.2014.00178.

Martín-Dacal, M., Fernández-Calvo, P., Jiménez-Sandoval, P., López, G., Garrido-Arandía, M., Rebaque, D., Hierro, I.d., Berlanga, D.J., Torres, M.Á., Kumar, V., et al. (2022). Arabidopsis immune responses triggered by cellulose and mixed-linked glucan oligosaccharides require a group of Leucine-Rich Repeat-Malectin Receptor Kinases. The Plant Journal 10.1111/tpj.16088.

Mergner, J., Frejno, M., List, M., Papacek, M., Chen, X., Chaudhary, A., Samaras, P., Richter, S., Shikata, H., Messerer, M., et al. (2020). Mass-spectrometry-based draft of the Arabidopsis proteome. Nature 579:409–414. 10.1038/s41586-020-2094-2.

Minh, B.Q., Schmidt, H.A., Chernomor, O., Schrempf, D., Woodhams, M.D., von Haeseler, A., and Lanfear, R. (2020). IQ-TREE 2: New Models and Efficient Methods for Phylogenetic Inference in the Genomic Era. Mol Biol Evol 37:1530–1534. 10.1093/molbev/msaa015.

Miya, A., Albert, P., Shinya, T., Desaki, Y., Ichimura, K., Shirasu, K., Narusaka, Y., Kawakami, N., Kaku, H., and Shibuya, N. (2007). CERK1, a LysM receptor kinase, is essential for chitin elicitor signaling in Arabidopsis. Proceedings of the National Academy of Sciences 104:19613–19618. 10.1073/pnas.0705147104.

Mohnen, D. (2008). Pectin structure and biosynthesis. Curr Opin Plant Biol 11:266–277. 10.1016/j.pbi.2008.03.006.

Molina, A., Jordá, L., Torres, M., Martín-Dacal, M., Berlanga, D.J., Fernández-Calvo, P., Gómez-Rubio, E., and Martín-Santamaría, S. (2024). Plant cell wall-mediated disease resistance: Current understanding and future perspectives. Mol Plant 17:699–724. 10.1016/j.molp.2024.04.003.

Moussu, S., Lee, H.K., Haas, K.T., Broyart, C., Rathgeb, U., De Bellis, D., Levasseur, T., Schoenaers, S., Fernandez, G.S., Grossniklaus, U., et al. (2023). Plant cell wall patterning and expansion mediated by protein-peptide-polysaccharide interaction. Science 382:719–725. 10.1126/science.adi4720.

Mélida, H., Sopeña-Torres, S., Bacete, L., Garrido-Arandia, M., Jordá, L., López, G., Muñoz-Barrios, A., Pacios, L.F., and Molina, A. (2018). Non-branched β-1,3-glucan oligosaccharides trigger immune responses in Arabidopsis. The Plant Journal 93:34–49. 10.1111/tpj.13755.

Mélida, H., Bacete, L., Ruprecht, C., Rebaque, D., Hierro, I.d., López, G., Brunner, F., Pfrengle, F., and Molina, A. (2020). Arabinoxylan-Oligosaccharides Act as Damage Associated Molecular Patterns in Plants Regulating Disease Resistance. Frontiers in Plant Science 11:1210. 10.3389/fpls.2020.01210.

Ngou, B.P.M., Heal, R., Wyler, M., Schmid, M.W., and Jones, J.D.G. (2022). Concerted expansion and contraction of immune receptor gene repertoires in plant genomes. Nature Plants 8:1146–1152. 10.1038/s41477-022-01260-5.

Nimchuk, Z.L., Zhou, Y., Tarr, P.T., Peterson, B.A., and Meyerowitz, E.M. (2015). Plant stem cell maintenance by transcriptional cross-regulation of related receptor kinases. Development 142:1043–1049. 10.1242/dev.119677.

Ochiai, A., Itoh, T., Kawamata, A., Hashimoto, W., and Murata, K. (2007). Plant Cell Wall Degradation by Saprophytic Bacillus subtilis Strains: Gene Clusters Responsible for Rhamnogalacturonan Depolymerization. Applied and Environmental Microbiology 73:3803–3813. 10.1128/aem.00147-07.

Parys, K., Colaianni, N.R., Lee, H.-S., Hohmann, U., Edelbacher, N., Trgovcevic, A., Blahovska, Z., Lee, D., Mechtler, A., Muhari-Portik, Z., et al. (2021). Signatures of antagonistic pleiotropy in a bacterial flagellin epitope. Cell Host & Microbe 29:620–634.e629. 10.1016/j.chom.2021.02.008.

Ralet, M.C., Tranquet, O., Poulain, D., Moïse, A., and Guillon, F. (2010). Monoclonal antibodies to rhamnogalacturonan I backbone. Planta 231:1373–1383. 10.1007/s00425-010-1116-y.

Ranf, S., Gisch, N., Schäffer, M., Illig, T., Westphal, L., Knirel, Y.A., Sánchez-Carballo, P.M., Zähringer, U., Hückelhoven, R., Lee, J., et al. (2015). A lectin S-domain receptor kinase mediates lipopolysaccharide sensing in Arabidopsis thaliana. Nature Immunology 16:426–433. 10.1038/ni.3124.

Richter, J., Watson, J.M., Stasnik, P., Borowska, M., Neuhold, J., Berger, M., Stolt-Bergner, P., Schoft, V., and Hauser, M.T. (2018). Multiplex mutagenesis of four clustered CrRLK1L with CRISPR/Cas9 exposes their growth regulatory roles in response to metal ions. Sci Rep 8:12182. 10.1038/s41598-018-30711-3.

Romanò, C., Awan, S., and Clausen, M.H. (2024). Chemical synthesis of branched rhamnogalacturonan I oligosaccharides. 10.26434/chemrxiv-2024-nlf7p.

Rozewicki, J., Li, S., Amada, K.M., Standley, D.M., and Katoh, K. (2019). MAFFT-DASH: integrated protein sequence and structural alignment. Nucleic Acids Res 47:W5–W10. 10.1093/nar/gkz342.

Ruprecht, C., Bartetzko, M.P., Senf, D., Dallabernadina, P., Boos, I., Andersen, M.C.F., Kotake, T., Knox, J.P., Hahn, M.G., Clausen, M.H., et al. (2017). A Synthetic Glycan Microarray Enables Epitope Mapping of Plant Cell Wall Glycan-Directed Antibodies. Plant Physiology 175:1094–1104. 10.1104/pp.17.00737.

Ruprecht, C., Bartetzko, M.P., Senf, D., Lakhina, A., Smith, P.J., Soto, M.J., Oh, H., Yang, J.Y., Chapla, D., Silva, D.V., et al. (2020). A Glycan Array-Based Assay for the Identification and Characterization of Plant Glycosyltransferases. Angewandte Chemie International Edition 59:12493–12498. 10.1002/anie.202003105.

Ryu, H., Choi, S., Cheng, M., Koo, B.K., Kim, E.Y., Lee, H.S., and Lee, D.H. (2025). Flagellin sensing, signaling, and immune responses in plants. Plant Commun:101383. 10.1016/j.xplc.2025.101383.

Schmidt, D., Schuhmacher, F., Geissner, A., Seeberger, P.H., and Pfrengle, F. (2015). Automated synthesis of arabinoxylan-oligosaccharides enables characterization of antibodies that recognize plant cell wall glycans. Chemistry 21:5709–5713. 10.1002/chem.201500065.

Schwessinger, B., Roux, M., Kadota, Y., Ntoukakis, V., Sklenar, J., Jones, A., and Zipfel, C. (2011). Phosphorylation-Dependent Differential Regulation of Plant Growth, Cell Death, and Innate Immunity by the Regulatory Receptor-Like Kinase BAK1. PLoS Genetics 7:e1002046. 10.1371/journal.pgen.1002046.

Senf, D., Ruprecht, C., de Kruijff, G.H., Simonetti, S.O., Schuhmacher, F., Seeberger, P.H., and Pfrengle, F. (2017). Active Site Mapping of Xylan-Deconstructing Enzymes with Arabinoxylan Oligosaccharides Produced by Automated Glycan Assembly. Chemistry 23:3197–3205. 10.1002/chem.201605902.

Smakowska-Luzan, E., Mott, G.A., Parys, K., Stegmann, M., Howton, T.C., Layeghifard, M., Neuhold, J., Lehner, A., Kong, J., Grunwald, K., et al. (2018). An extracellular network of Arabidopsis leucine-rich repeat receptor kinases. Nature 553:342–346. 10.1038/nature25184.

Tseng, Y.H., Scholz, S.S., Fliegmann, J., Krüger, T., Gandhi, A., Furch, A.C.U., Kniemeyer, O., Brakhage, A.A., and Oelmüller, R. (2022). CORK1, A LRR-Malectin Receptor Kinase, Is Required for Cellooligomer-Induced Responses in. Cells 1110.3390/cells11192960.

Voxeur, A., Habrylo, O., Guénin, S., Miart, F., Soulié, M.-C., Rihouey, C., Pau-Roblot, C., Domon, J.-M., Gutierrez, L., Pelloux, J., et al. (2019). Oligogalacturonide production upon Arabidopsis thaliana–Botrytis cinerea interaction. Proceedings of the National Academy of Sciences 116:19743–19752. 10.1073/pnas.1900317116.

Wang, L., Einig, E., Almeida-Trapp, M., Albert, M., Fliegmann, J., Mithöfer, A., Kalbacher, H., and Felix, G. (2018). The systemin receptor SYR1 enhances resistance of tomato against herbivorous insects. Nature Plants 4:152–156. 10.1038/s41477-018-0106-0.

Wanke, A., Rovenich, H., Schwanke, F., Velte, S., Becker, S., Hehemann, J.H., Wawra, S., and Zuccaro, A. (2020). Plant species-specific recognition of long and short β-1,3-linked glucans is mediated by different receptor systems. The Plant Journal 102:1142–1156. 10.1111/tpj.14688.

Willats, W.G., McCartney, L., Mackie, W., and Knox, J.P. (2001). Pectin: cell biology and prospects for functional analysis. Plant Mol Biol 47:9–27.

Wolf, S. (2022). Cell Wall Signaling in Plant Development and Defense. Annu Rev Plant Biol 73:323–353. 10.1146/annurev-arplant-102820-095312.

Wu, Y., Xun, Q., Guo, Y., Zhang, J., Cheng, K., Shi, T., He, K., Hou, S., Gou, X., and Li, J. (2016). Genome-Wide Expression Pattern Analyses of the Arabidopsis Leucine-Rich Repeat Receptor-Like Kinases. Molecular Plant 9:289–300. 10.1016/j.molp.2015.12.011.

Xiao, Y., Sun, G., Yu, Q., Gao, T., Zhu, Q., Wang, R., Huang, S., Han, Z., Cervone, F., Yin, H., et al. (2024). A plant mechanism of hijacking pathogen virulence factors to trigger innate immunity. Science 383:732–739. 10.1126/science.adj9529.

Yamaguchi, Y., Huffaker, A., Bryan, A.C., Tax, F.E., and Ryan, C.A. (2010). PEPR2 is a second receptor for the Pep1 and Pep2 peptides and contributes to defense responses in Arabidopsis. Plant Cell 22:508–522. 10.1105/tpc.109.068874.

Zakharova, A.N., Madsen, R., and Clausen, M.H. (2013). Synthesis of a backbone hexasaccharide fragment of the pectic polysaccharide rhamnogalacturonan I. Org Lett 15:1826–1829. 10.1021/ol400430p.

